# Key Genetic Determinants Driving Esophageal Squamous Cell Carcinoma Initiation and Immune Evasion

**DOI:** 10.1101/2022.10.13.512143

**Authors:** Kyung-Pil Ko, Yuanjian Huang, Shengzhe Zhang, Gengyi Zou, Bongjun Kim, Jie Zhang, Sohee Jun, Cecilia Martin, Karen J. Dunbar, Gizem Efe, Anil K. Rustgi, Hiroshi Nakagawa, Jae-Il Park

## Abstract

**Background and aims:** Despite recent progress in identifying aberrant genetic and epigenetic alterations in esophageal squamous cell carcinoma (ESCC), the mechanism of ESCC initiation remains unknown.

**Methods:** Using CRISPR/Cas 9-based genetic ablation, we targeted 9 genes (*TP53*, *CDKN2A*, *NOTCH1*, *NOTCH3*, *KMT2D*, *KMT2C*, *FAT1*, *FAT4*, and *AJUBA*) in murine esophageal organoids (EOs). Transcriptomic phenotypes of organoids and chemokine released by organoids were analyzed by single-cell RNA sequencing (scRNA-seq). Tumorigenicity and immune evasion of organoids were monitored by allograft transplantation. Human ESCC scRNA-seq datasets were analyzed to classify patients and find subsets relevant to organoid models and immune evasion.

**Results:** We established 32 genetically engineered EOs and identified key genetic determinants that drive ESCC initiation. A single-cell transcriptomic analysis uncovered that *Trp53*, *Cdkn2a*, and *Notch1* (PCN) triple-knockout (KO) induces neoplastic features of ESCC by generating cell lineage heterogeneity and high cell plasticity. *PCN* KO also generates an immunosuppressive niche enriched with exhausted T cells and M2 macrophages via the CCL2-CCR2 axis. Mechanistically, *CDKN2A* inactivation transactivates *CCL2* via NF-κB. Moreover, comparative single-cell transcriptomic analyses stratified ESCC patients and identified a specific subtype recapitulating the PCN-type ESCC signatures, including the high expression of CCL2 and CD274/PD-L1.

**Conclusions:** Our study unveils that loss of *TP53*, *CDKN2A*, and *NOTCH1* induces esophageal neoplasia and immune evasion for ESCC initiation and proposes the CCL2 blockade as a viable option for targeting PCN-type ESCC.

## Introduction

ESCC accounts for over 80% of all esophageal cancer cases and has a poor prognosis because of a lack of symptoms in the early stages. The 5-year overall survival rate of patients with esophageal cancer ranges from 10% to 25%.^1^ ESCC develops from squamous dysplasia and is asymptomatic until it spreads to other tissues,^2^ which leads to late detection. Therefore, understanding the mechanisms of esophageal neoplasia and ESCC initiation is imperative to improving the detection, diagnosis, treatment, and prevention of ESCC. However, despite a recent genome-wide analysis of ESCC patients, the key genetic factors that drive ESCC neoplasia and initiation remain elusive.

In addition to the genetic and epigenetic alteration of tumor cells, the tumor microenvironment (TME)^3^ plays a pivotal role in tumorigenesis. The TME comprising immune and stromal cells surrounding tumor cells designs the immune landscape to elicit immunosuppressive effects on tumors and various immunotherapeutic responses.^4, 5^ Although the TME’s effects on tumor cells are biphasic, as immune cells generally recognize tumor cells or neoantigens to generate anti-tumor immune responses,^6^ the tumor-favorable TME induces tumor growth, invasion, and metastasis.^7^ The TME is also an indispensable factor in ESCC development, as it provides drug resistance and immunosuppressive immune cell infiltration.^8, 9^

Immune checkpoint inhibitors (ICIs) have been clinically tested in ESCC patients and have shown promising results.^10–14^ However, not all patients benefit from ICIs, alone or in combination with first-line therapy,^8^ possibly because of the lack of a clear classification of ESCC patients. The underlying genetic background of immunosuppressive microenvironment has yet to be identified. Herein, we found that the genetic inactivation of *TP53*, *CDKN2A*, and *NOTCH1* induces esophageal neoplasia (cell-autonomous) and generates an immunosuppressive niche (non-cell-autonomous) to promote ESCC tumorigenesis.

## Methods

### Mice

C57BL/6, *Trp53^floxed/floxed^* (JAX no. 008462) and *Rosa26nT-nG* (JAX no. 023035) mice purchased from the Jackson Laboratory were bred and housed in the Animal facility at MD Anderson Cancer Center on a 12-hr light/dark cycle. All animal procedures were performed according to the guidelines of the Association for the Assessment and Accreditation of Laboratory Animal Care and institutionally approved protocols. The study was compliant with all relevant ethical regulations regarding animal research.

### Wnt3A, R-Spondin1, and Noggin-Conditioned Medium

The Wnt3A, R-Spondin1, and Noggin (WRN)-conditioned medium was prepared as previously described.^15^ In brief, L-WRN (ATCC CRL-3276) cells were cultured on a 10-cm plate with culture medium (Dulbecco’s modified Eagle’s medium [DMEM, Fisher], 0.5 mg/mL G418 (Thermofisher), 0.5 mg/mL hygromycin B (Thermofisher), 1% penicillin/streptomycin (Life Technologies), and 10% fetal bovine serum). After 10% of L-WRL cells (ATCC, CRL-3276) had been seeded in culture medium (without G418 and hygromycin B) in 10-cm plates, cells were incubated for 3-4 days. The medium was replaced with 10 mL of fresh medium when the cells were 80%-90% confluent, and the cells were incubated for 24 hrs. The medium was collected, centrifuged at 1000 × g for 4 min, passed through a 0.22-μm sterile filter, and stored at −80 °C. Another 10 mL of fresh medium was added to the plates and collected after 24 hrs to make the second batch of conditioned medium using the same procedures. The first, second, and third batches of conditioned media were mixed before use to prepare a 100% WRN-conditioned medium.

### Esophageal Tissue Isolation

Tissues were collected from mice (8∼10-week-old) euthanized via CO_2_ inhalation followed by cervical dislocation. The esophagi were collected in 10-cm Petri dishes with ice-cold phosphate-buffered saline (PBS) with 1% penicillin/streptomycin and swirled gently to remove blood. The esophagi were opened longitudinally and washed with cold PBS. The epithelial cell layer was peeled off using surgical tweezers and then dissected into 0.5-cm^3^ pieces with surgical blades. The minced esophagi were collected in a 15-mL conical tube and digested by 0.05% trypsin-EDTA (Thermofisher) at 37 L for 60 min with frequent vortexing. After dissociation, 3× volume 10% fetal bovine serum-supplemented DMEM was added, followed by vigorous pipetting. The suspension was passed through a 35-μm cell strainer to collect a single-cell suspension. Finally, the cell suspension was spun down at 1000 rpm for 4 min and resuspended in a 50% WRN-conditioned medium.

### EO Culture

The E-MEOM was used to culture EOs. Single-cell–dissociated esophageal epithelial cells (1000 cells) were suspended in 8 μL of E-MEOM and 12 μL of pre-thawed Matrigel (Corning) on ice and seeded in the centers of a well to create a Matrigel dome. The plate was incubated at 37 °C for 10 min to solidify the Matrigel. Finally, 500 μL of E-MEOM was added and incubated at 37 °C with 5% CO_2_. The medium was changed every 2-3 days.

### Trp53KO EO

Esophagi from 8-to 10-week-old *Trp53^floxed/floxed^* mice were digested into a single-cell suspension and seeded in Matrigel to form EOs, as described above. After 7 days of culture, EOs were digested with 0.05% trypsin-EDTA at 37 °C for 45 min to create a single-cell suspension. Ad-CMV-EGFP or Ad-Cre-EGFP (University of Iowa) was added to the cell suspension at 1 × 10^3^ pfu/cell. Cells were then suspended in Matrigel to generate new EOs. After 2 days of seeding, 10 μM nutlin3 was added for *Trp53^floxed/floxed^*cell selection. Two days later, the selection was performed by sorting the GFP+ cells after dissociating EOs into single cells, and the sorted cells were re-seeded for EO culture, as described above. EOs were collected for genotyping 7 days after seeding. Wild-type, *Trp53^floxed/floxed^*, and KO Trp53 alleles were amplified as 288 bp, 370 bp, and 612 bp, respectively. See Supplementary Table 1 for primer information.

### CRISPR/Cas9-Based Gene Targeting in EOs

CRISPR/Cas9 system-mediated gene KO was described in a previous study.^16^ In brief, WT or *Trp53* KO EOs were digested with 0.05% trypsin-EDTA to dissociate into single cells. Single cells were incubated with a virus-containing medium with polybrene for 1 hr with centrifugation (600 g) at 32 °C (see Supplementary Table 2 for sgRNA sequences). Cells were then incubated at 37 °C with 5% CO_2_ for 4 more hrs and embedded in Matrigel. The medium was replaced 2 days after infection with antibiotics (puromycin [Sigma], blasticidin [Invitrogen], or hygromycin) for selection. Gene KO was confirmed by genomic DNA PCR (see Supplementary Table 1 for primer information). Additional details are described in the Supplementary Methods.

## Results

### Genetic ablation of *Trp53* and *Notch1* is sufficient to induce esophageal neoplasia

To elucidate the vital genetic events initiating ESCC, we analyzed genetic alterations in ESCC patients (*n* = 86). Recurrent loss-of-function mutations in *TP53*, *CDKN2A*, *KMT2D*, *KMT2C*, *FAT1*, *FAT4*, *AJUBA*, *NOTCH1*, and *NOTCH3* genes were observed in ESCC patients.^17, 18^ The *TP53* and *CDKN2A* genes were the most commonly altered (> 70%) (Figure S1A). More than 50% of mutations in the *CDKN2A*, *KMT2D*, *NOTCH1*, and *FAT1* genes were frameshift or truncation mutations (Figure S1B). Additionally, ESCC patients displayed transcriptional downregulation of *KMT2C* and *FAT4* genes (Figure S1C). To understand the significance of the nine genes (*TP53*, *CDKN2A*, *KMT2D*, *KMT2C*, *NOTCH1*, *NOTCH3*, *FAT1*, *FAT4*, and *AJUBA*) in ESCC initiation, we tested the functional impact of their inactivation on the transformation of esophageal organoids (EOs), as recently established (Figure 1A, S2A).^19, 20^ Employing the CRISPR system, we genetically ablated the nine genes in 32 different combinations. Given that *TP53* and *CDKN2A* genes were the most frequently mutated, *Trp53* KO and *Cdkn2a* KO were chosen as the foundation for additional gene ablations, and double KO (dKO) or triple KO (tKO) EOs were established (Figure S1D-F, S2B,C).

**Figure 1.**
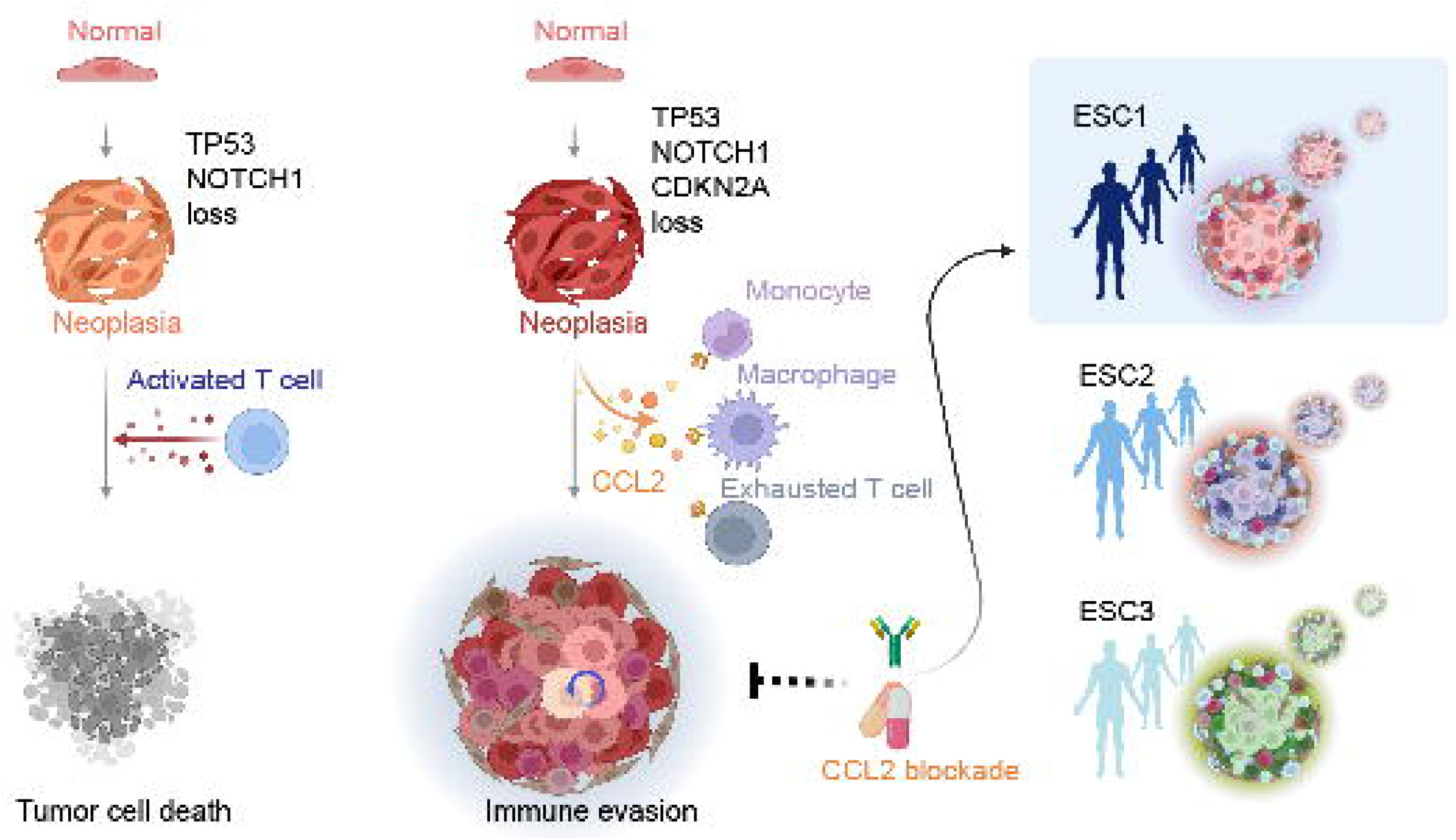
Genetic ablation of *Trp53* and *Notch1* induces esophageal hyperplasia and de-differentiation. **A,** Schematic structure of murine EOs. **B,** Heatmap visualizing the volume of each EO. **C,** Hematoxylin-and-eosin (H&E) images; scale bars = 50 μm. **D,E,** Immunofluorescent (IF) images of EOs (D); MKI67+ cell quantification (E). WT: *Trp53^floxed/floxed^*, N: *Notch1* KO, C: *Cdkn2a* KO, CN: *Cdkn2a* KO + *Notch1* KO, P: *Trp53^del/del^*, PN: *Trp53^del/del^*+ *Notch1* KO, PC: *Trp53^del/del^* + *Cdkn2a* KO, PCN: *Trp53^del/del^* + *Cdkn2a* KO + *Notch1* KO; scale bars = 20 μm; \*\*\*\**P* < 0.0001. **F,** IF staining images; scale bars = 20 μm. **G,H,** BrdU incorporation assays; IF with anti-BrdU antibody (G); BrdU+ cell quantification (H); scale bars = 10 μm; \*\*\*\**P* < 0.0001. **I,J,** Bright-field images of EOs co-cultured with mesenchymal (Tomato [red] fluorescent-expressing) cells (I); E-cadherin/CDH1 IF images of EOs (J); scale bars = 50 μm. **K,L,** Assessing colony formation ability (K) and cell growth rate (L) of EOs.

Combined with *Trp53* and *Cdkn2a* dKO, genetic ablation of *Notch1*, *Fat4*, or *Ajuba* resulted in markedly increased EO size (Figure 1B). However, only *Notch1* KO with *Trp53* and *Cdkn2a* dKO EOs displayed cell differentiation (central keratin pulp) loss and hyperplastic growth simultaneously (Figure 1B and C). KO of *Ajuba* or *Fat4* with *Trp53* or *Cdkn2a* dKO was insufficient to induce neoplastic transformation, as these cells showed keratin pulp (well-differentiated). *Notch1* KO induced loss of keratin layer at the center of organoids, regardless of *Trp53* and *Cdkn2a* status, indicating a major role of NOTCH signaling in esophageal epithelial cell differentiation (Figure 1C). The increased number of proliferating cells was confirmed by MKI67 staining in dKO of *Trp53* and *Notch1* (PN) and tKO of *Trp53*, *Cdkn2a*, and *Notch1* (PCN) EOs (Figure 1D,E, S3A). Compared to other EOs, PN and PCN EOs showed upregulated expression of ESCC stemness markers (Sox2 and Trp63) and increased cell proliferation (Figure 1D-H, S3A). *Notch1* KO suppressed organoid cell differentiation, as represented by decreased Krt13, a cell differentiation marker (Figure 1F, S3B-D).

Since ESCC invades stromal cell layers at an early stage,^21, 22^ we analyzed EO cell invasiveness by co-culturing EOs on the stromal cell layer. While WT EO-derived epithelial cells formed distinct epithelial cell colonies, PN and PCN EO-derived cells were mixed with a stromal cell layer (Figure 1I,J, S3E), recapitulating ESCC’s invasive feature. We further examined the tumorigenicity of each group of EOs in 2D culture by assessing cell growth without supporting materials and growth factors in the organoid culture system. In 2D culture, only PN and PCN EOs showed increased colony formation, exponential cell growth, and cell migration, whereas PCA and PCF4 organoids showing increased size in 3D culture did not proliferate without the supporting factors (Figure S3H). Notably, PN EOs showed relatively faster growth than did PCN EOs (Figure 1K,L, S3F,G). Additionally, AB-PAS staining showed that PN and PCN EOs did not express mucins highly expressed in esophageal adenocarcinoma (EAC), indicating that PN- and PCN-induced neoplasia is unrelated to EAC (Figure S3I). Furthermore, since the upper esophageal cells were the source of the organoids, it is inherently unlikely that EAC development would occur. These results suggest that *Trp53* and *Notch1* dKO (PN) and *Trp53*, *Cdkn2a*, and *Notch1* tKO (PCN) are sufficient to induce neoplastic transformation that exhibits the characteristics of early-stage ESCC.

### Dysregulated cell plasticity and lineage trajectories of neoplastic EOs

Next, to understand the mechanism of ESCC initiation, we performed multiplexed single-cell RNA-sequencing (scRNA-seq) of WT, PC, PN, and PCN EOs (Figure 2A-C, S4A). Thirty-five cell clusters were annotated with multiple markers that are specifically expressed in proliferative (*Mki67*, *Top2a*, and *Stmn1*), early suprabasal (*Ptma*, *Itgb4*, and *Krt5*), intermediate suprabasal (*Itga6*), and stratified (*Krt13*, *Krt4*, and *Sprr3*) cell types (Figure S4B-D).^19, 23^ The intermediate suprabasal cell cluster showed a moderate expression level of early suprabasal markers (*Itgb4* and *Krt5*) and stratified markers. The proliferative cells were significantly enriched in the PC, PN, and PCN organoids compared to in the WT EOs mainly composed of differentiated (stratified and suprabasal) cells (Figure 2D, S4E), recapitulating EO phenotypes (Figure 1).

**Figure 2.**
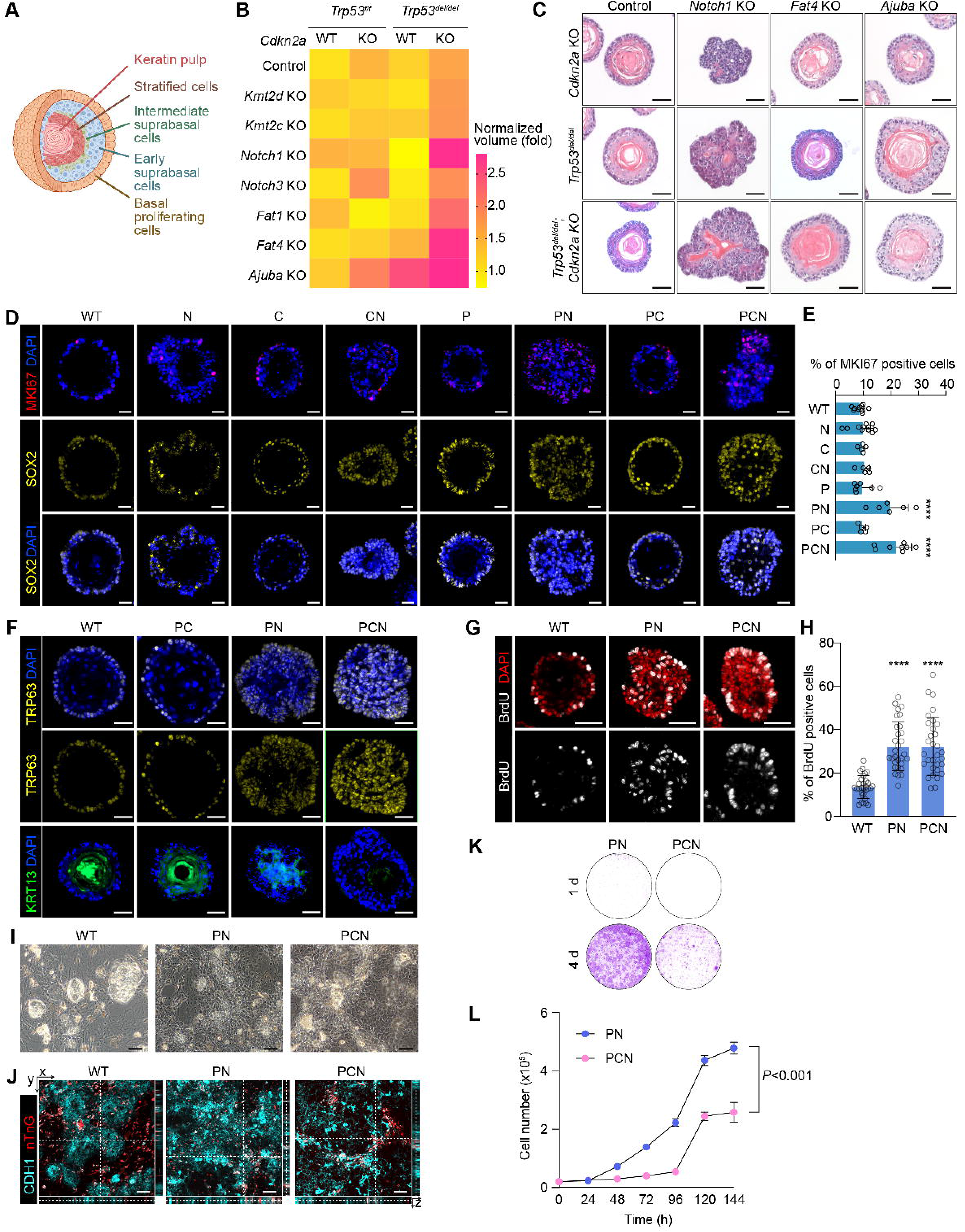
Single-cell transcriptomic analysis of genetically engineered EOs. **A,** Schematic overview of scRNA-seq procedure. Four types of EOs (WT, PC, PN, and PCN) were multiplexed for library preparation. **B,C,** UMAPs of four integrated datasets. **D,** The proportion of each cluster was compared to the same cluster of the WT dataset. Statistical significance between 2 groups was assessed by the permutation test. **E,** RNA velocity–based UMAP projection based on cell types in each dataset. **F,** Cell trajectory inference by RNA velocity. **G,** Latent time results projected on the velocity-based UMAP. **H,** PAGA analysis of RNA velocity–based cell clusters showing the direction of cell lineage on UMAP. The size of the circle corresponds to the cell number. **I,** Representative marker genes of root cell clusters with simplified lineage trajectories.

Having observed distinct cell hyperproliferation and de-differentiation by PN and PCN, we performed RNA velocity-based cell lineage trajectory inference (Figure 2E). Intriguingly, PN and PCN showed multiple root cell clusters (PN3, PN4, PN8 and PCN3, PCN4, and PCN7, respectively), whereas WT harbored a single root cell cluster (WT4), also reproduced in the latent time and PAGA analyses (Figure 2F-I). These results imply that *Trp53* and *Notch1* dKO might generate distinct cellular plasticity and higher cellular heterogeneity.^24^

### Human ESCC features in *Trp53*/*Notch1* dKO and *Trp53*/*Cdkn2a*/*Notch1* tKO EOs

We assessed the pathological relevance of genetically engineered EOs to human ESCC by comparing their transcriptional signatures. Compared to WT and PC EOs, ESCC cancer stem cell markers, including *Trp63*, *Krt5*, and *Tfrc*, were intensively enriched in the proliferating cells of PN and PCN EOs (Figure 3A). *Sox2* and *Gnl3* were highly expressed in the proliferating cells of PN and PCN EOs, respectively, while the other genes did not show specificity to the PN or PCN cell types. We also compared the transcriptional features of our EOs with those of ESCC patients from the TCGA database using the Scissor (v2.0.0) package. The gene expression feature comparison analysis showed the similarity of each EO to ESCC patients: WT (35.4%), PC (52.2%), PN (50.5%), and PCN (52.2%) (Figure 3B,C). In addition, PCN (21.9%) showed the highest similarity with the poor survival–associated ESCC patients compared to the other organoids (Figure 3D,E). A gene set enrichment analysis further confirmed that the PCN-highly expressed gene set was enriched in ESCC-specific gene features, while the PN-highly expressed gene set enrichment was not statistically significant (Figure 3F). To increase the sequencing read depth, we also performed bulk RNA-seq of WT, PN, and PCN EOs and reanalyzed the gene expression of each dataset (Figure 3G). Using the differentially expressed genes (DEGs) of PN to WT or PCN to WT, we ran the Enrichr analysis with PN or PCN highly expressed genes. Both PN and PCN highly expressed genes were enriched with features of the cell cycle, DNA replication, and mitotic cells in the Bioplanet and REACTOME databases (Figure 3H). The “pathways in cancer” were also associated with the PCN highly expressed genes in the Kyoto Encyclopedia of Genes and Genomes (KEGG) pathway enrichment analysis (Figure 3I). These results suggest that the transcriptomic patterns of human ESCC are most similar to the transcriptomic features of *Trp53*, *Cdkn2a*, and *Notch1* tKO EOs.

**Figure 3.**
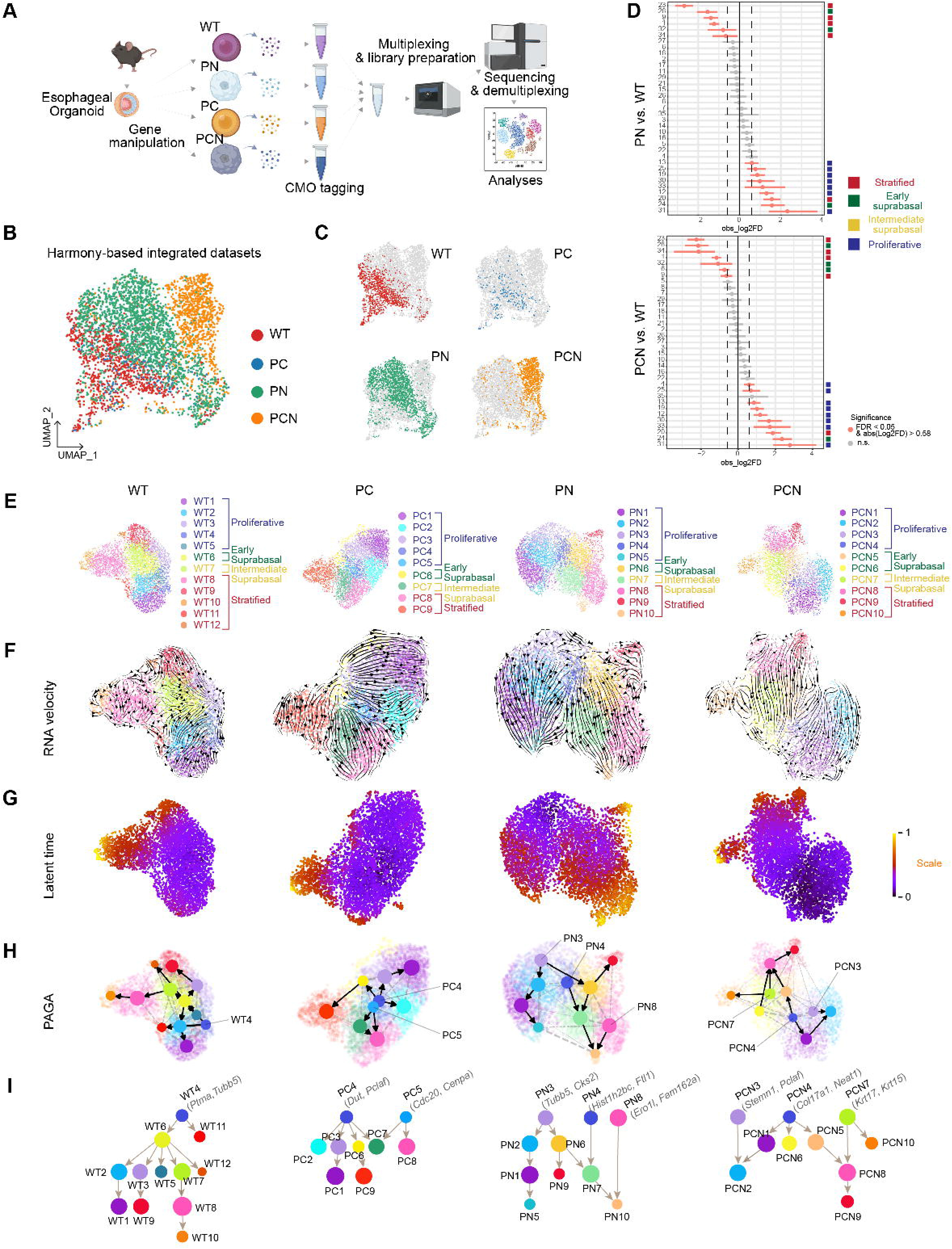
Transcriptomes of PN and PCN recapitulate the ESCC phenotype. **A,** Dot plot of ESCC stem cell markers. **B,C,** ESCC phenotype–associated cells were marked as ESCC^+^ cells in Scissor-based UMAP projection (B), and the proportion of ESCC^+^ cells is shown in bar plot (C). \*\*\*\**P* value < 0.0001, as determined by Fisher’s exact test. **D,E,** Poor survival of ESCC patient–associated cells is displayed as Poor survival^+^ cells in UMAP (D), and the proportion was analyzed (E). \*\*\*\**P* value < 0.0001, as determined by Fisher’s exact test. n.s. = not significant. **F,** Gene set enrichment analysis of PN or PCN dataset. **G,** Heatmap clustering of bulk RNA-sequencing from WT, PN, and PCN EOs. **H,** Enrichr from bulk RNA-seq datasets. PN- or PCN-enriched genes compared to WT were analyzed using Bioplanet and REACTOME databases; TOP5 features of each database are shown; \**P* value. **I,** Dot plots of PN or PCN highly expressed genes compared to gene sets of the Kyoto Encyclopedia of Genes and Genomes (KEGG) pathways; dot size: gene number count; dot colors: adjusted *P* values.

### *Trp53*/*Cdkn2a*/*Notch1* tKO cells are tumorigenic *in vivo* via CCL2-mediated immune evasion

Having determined that either PN dKO or PCN tKO is sufficient to induce the neoplastic phenotype *in vitro* (Figures 1, 2), we determined their impact on ESCC tumorigenesis *in vivo* by congenic transplantation. Subcutaneously injected PN and PCN cells developed tumors in immunocompromised mice (Figure S5A-E), consistent with the *in vitro* results. However, in immunocompetent mice, only PCN cells exhibited tumor growth, with a success rate of 60%, whereas PN cells failed to proliferate (0 of 5) (Figure 4A,B). This finding was intriguing, as it was unexpected given the superior growth rate of PN cells compared to that of PCN cells *in vitro* (Figure 1K,L). PCN cell–derived tumors had substantially invaded the muscular layer with poorly differentiated squamous cell carcinoma (Figure 4B). PCN tumors were hyperproliferative, with ESCC stemness marker (SOX2, TRP63) expression, recapitulating the features of ESCC (Figure 4C).

**Figure 4.**
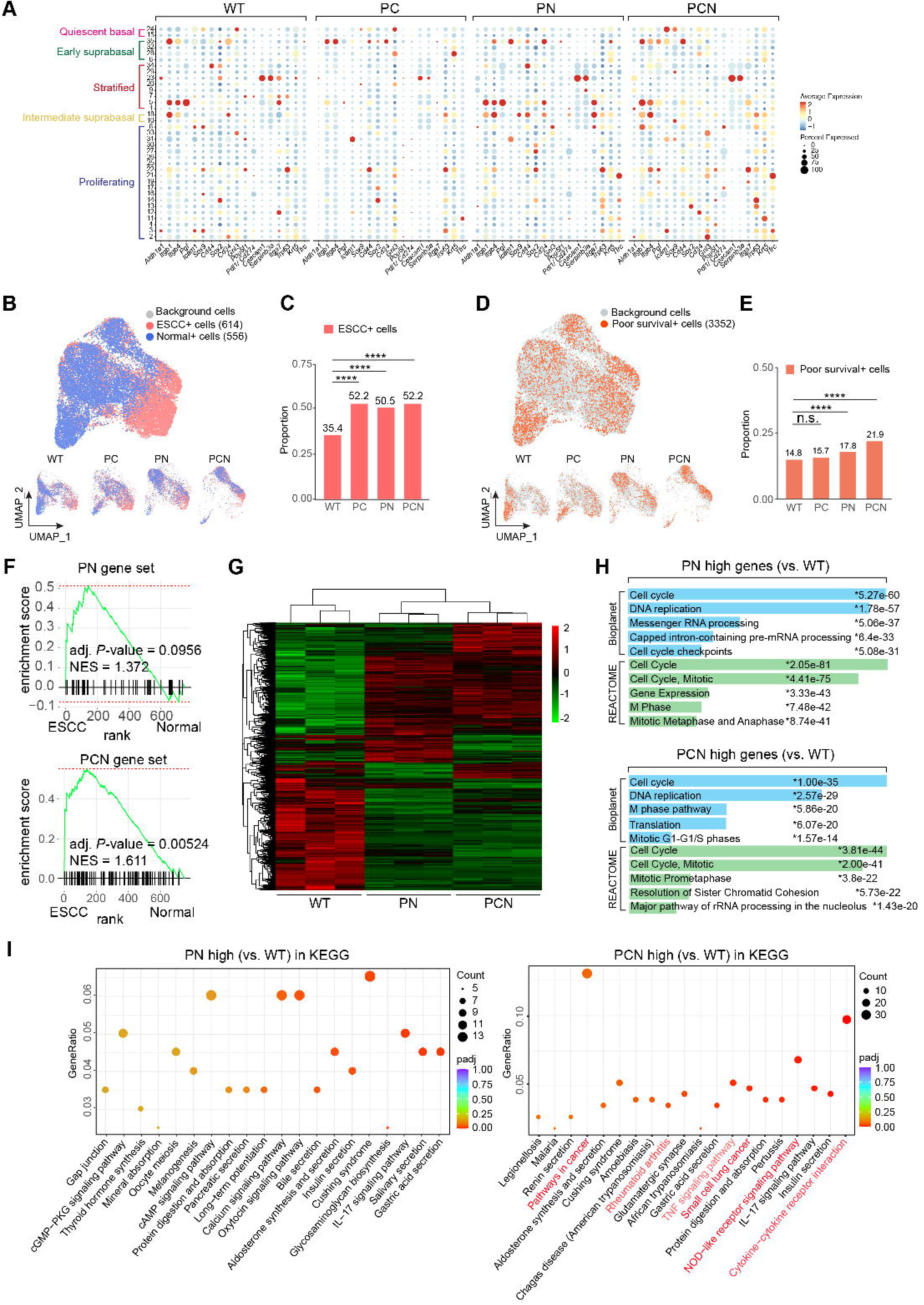
*In vivo* tumorigenicity of PCN with pro-tumorigenic TME. **A,** Assessment of tumor growth from C57BL/6 mice subcutaneously injected with either PN or PCN cells. **B,** H&E images; red arrowheads, mitotic cells; black arrowhead, blood vessel; dotted circle, inflammatory cells; scale bars = 100 μm. **C,** IF images of transplanted tumors; scale bars = 100 μm. **D,** Enrichr analysis from DEGs of PCN and PN scRNA-seq datasets. **E,** Volcano plot of DEGs between PCN and PN datasets. **F,** Feature plot of *Ccl2*. **G-I,** Tumor assessment of mice subcutaneously transplanted with PCN or PCN-*Ccl2* KO cells; tumor images (G), tumor volume (H), and weight (I) comparison. **J,** PDCD1^+^, PRF1/PERFORIN^+^, CD8^+^, CD206^+^, CD209^+^, and MKI67^+^ cells in randomly chosen 630× magnified images were analyzed and plotted. n.s. = not significant. **K,** PDCD1/PD-1, PRF1/PERFORIN, CD8, CD206, CD209, and MKI67 staining of PCN and PCN-*Ccl2* KO–derived tumors. Scale bars = 20 μm.

These results led us to hypothesize that PCN cells evade immune surveillance, while PN cells do not. Indeed, CD4^+^ T cells, CD8^+^ T cells, neutrophils, and dendritic cells infiltrated the PCN-derived tumors, implying an active interaction between tumor and immune cells (Figure S5F). To test this, we comparatively examined PN and PCN transcriptomes. An Enrichr analysis showed that the genes associated with T-cell receptor signaling and antigen processing and presentation were highly enriched in the PCN compared to in the PN (Figure S5G). A DEG analysis of bulk RNA-seq showed that the genes related to immune response and cytokine-related signaling pathways were highly expressed in PCN compared to in PN (Figure 4D, S5H), suggesting that cytokines or chemokines specifically expressed in PCN cells contribute to immune evasion. A comparative DEG analysis of scRNA-seq datasets identified three cytokines (*Ccl2*, *Cxcl1*, and *Cxcl2*) highly upregulated in PCN (Figure 4E). Notably, only *Ccl2* expression was enriched explicitly in the PCN compared to in WT, PC, and PN (Figure 4F, S5I-M), validated by immunostaining of PCN allograft tumors for CCL2 and its receptor, CCR2 (Figure S5N).

Since CCL2 expressed by tumor cells recruits myeloid cells (monocytes, tumor-associated macrophages [TAMs], and myeloid-derived suppressor cells [MDSCs]) to the TME, inhibiting the CCL2-CCR2 axis was shown to enhance the immune response to tumors.^25–28^ To determine the role of Ccl2 in immune evasion of PCN cells, we genetically ablated the *Ccl2* gene in PCN cells (PCN-*Ccl2* KO) and compared the tumorigenicity of PCN and PCN-*Ccl2* KO cells using allograft transplantation. PCN and PCN-*Ccl2* KO showed no difference in cell growth *in vitro* (Figure S6A,B) or tumor growth in immunocompromised mice (Figure S6C,D). However, unlike PCN cells, PCN-*Ccl2* KO cells barely developed tumors in immunocompetent mice (Figure 4G-I). Although immune cells were abundant in both PCN parental and *Ccl2* KO tumors, the keratin pearls, one of the signatures of ESCC, were mainly observed in PCN tumors compared to in *Ccl2* KO tumors, indicating that PCN tumors develop with less interruption by immune cells (Figure S6E). In line with this, the number of exhausted T (T_ex_) cells (expressing Pdcd1/Pd-1) was increased among the immune cells surrounding PCN tumors compared to those surrounding PCN-*Ccl2* KO tumors (Figure 4J,K, S6F). Moreover, PCN-*Ccl2* KO tumors showed a higher number of cells with perforin, an effector CD8^+^ T cell marker, CD8^+^ T cells, and cleaved-Caspase-3^+^ (an apoptotic cell marker) cells compared to PCN tumors, suggesting that more cell death mediated by T cells occurs in PCN-*Ccl2* KO tumors than in PCN tumors (Figure 4J,K, S6G).

Simultaneously, a more significant number of CD206^+^ (an M2 macrophage marker) cells was observed in PCN tumors than in PCN-*Ccl2* KO tumors (Figure 4J,K, S6F).

Although we found fewer M2 macrophage marker–expressing cells in PCN-*Ccl2* KO tumors than in PCN tumors, we did not find significant differences in the number of infiltrating M1 macrophages (CD68^+^/CD80^+^ cells) or MDSCs (CD11B^+^/LY6G^+^ cells) (Figure S6F-H). Nonetheless, the proliferation of PCN and PCN-*Ccl2* KO tumors was comparable based on the MKI67^+^ cell quantification (Figure 4J,K, S6F). These results suggest that CCL2 secreted from PCN tKO tumor cells contributes to immune evasion in ESCC tumorigenesis.

### Emerging CCL2-CCR2 interaction between tumor and immune cells during ESCC tumorigenesis

To complement allograft models, we analyzed single-cell transcriptomics of an autochthonous mouse model.^29^ In this model, treatment of 4-NQO (4-nitroquinoline 1-oxide) resulted in the development of inflammation, hyperplasia, dysplasia, cancer in situ, and invasive cancer in the esophageal epithelium in a time-dependent manner. From the scRNA-seq datasets of the 4-NQO model, the epithelial cells (proliferating, basal, suprabasal, stratified, and other cells) and the immune cells were sub-clustered based on marker gene expression (Figure S7A-F). This ESCC model showed decreased expression of Trp53, Cdkn2a, and Notch signaling pathway-related genes through ESCC development, similar to PCN organoids (Figure S8A). A comparative analysis of epithelial cell transcriptomes at each stage in ESCC patient tumors showed increasing similarity as murine ESCC develops under 4-NQO exposure (Figure S8B,C), suggesting that the 4-NQO mouse model recapitulates human ESCC tumorigenesis. Notably, *Ccl2* and *Ccr2* expression in the esophageal epithelial and immune cells, respectively, was the highest in hyperplasia (Figure 5A,B, S8D,E). A further subtype analysis showed that *Ccr2* expression was enriched in the T_ex_ cells and regulatory T (T_reg_) cells (Figure S8F). Moreover, we found that T_ex_ cell marker (*Cd160*, *Havcr2*/*Tim-3*, *Pdcd1*/*Pd-1*, and *Lag3*) expression was increased in the T cells at the early stage of ESCC tumorigenesis (Figure S8G), consistent with our PCN-transplanted tumor cells (Figure 4, S6). To investigate the intercellular CCL-mediated ligand-receptor interaction, epithelial and immune cell communications were analyzed using the Squidpy and CellChat packages. The CCL2-CCR2 interaction was inferred to be strong from the hyperplasia state and maintained until invasive cancer (Figure S8H). Moreover, the total interactions related to the CCL pathway between epithelial cells and immune cells were increased during ESCC development (Figure 5C, S8I). The CCL pathway–specified cell-cell communications were observed starting from the inflammation status in epithelial cells, T_ex_ cells, MDSCs, and macrophages (Figure 5D, S8J).

**Figure 5.**
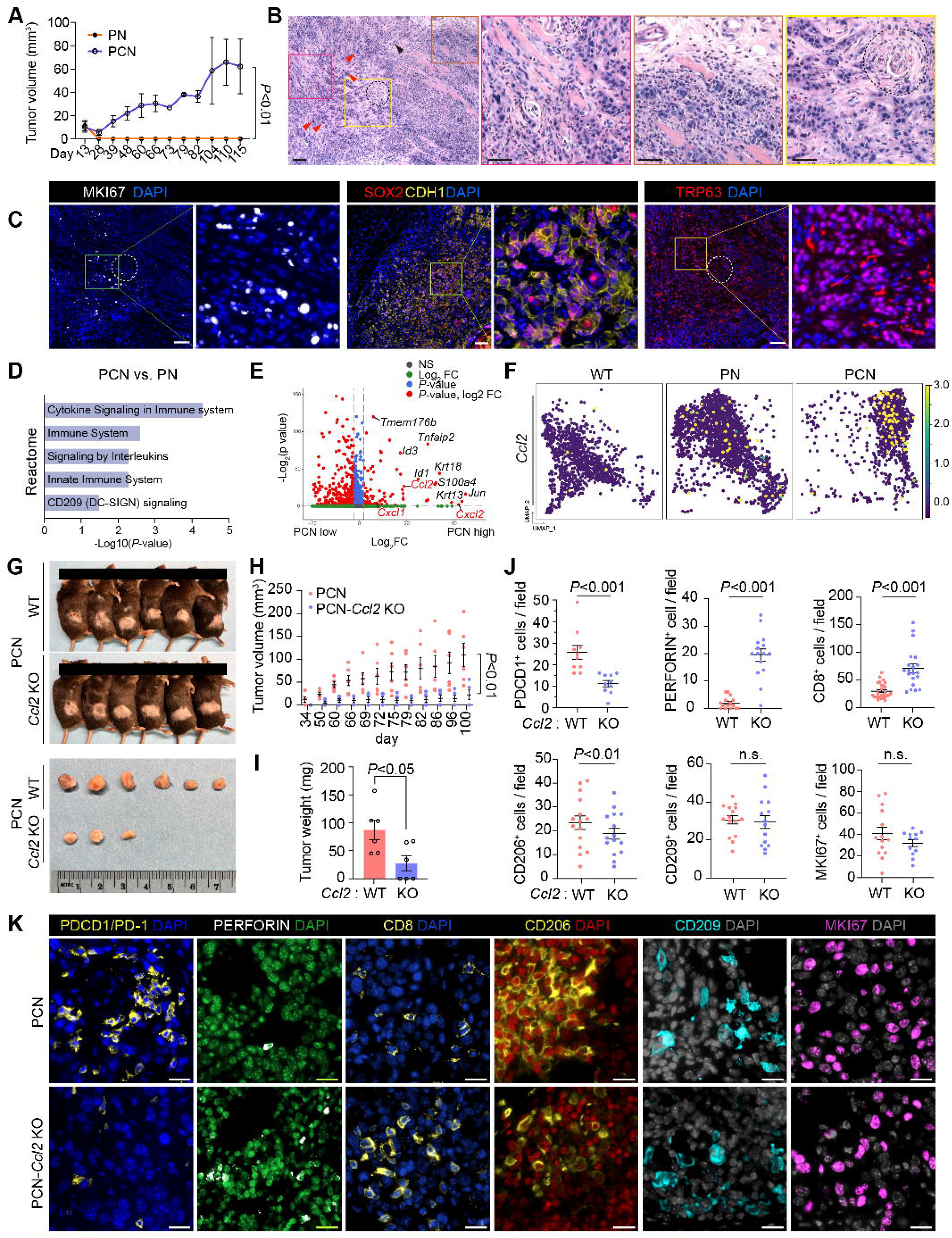
CCL2-CCR2-induced immune evasion during ESCC development. **A,** Dot plot visualizing epithelial cells’ *Ccl2* expression of each disease status. **B,** Dot plot showing *Ccr2* expression in immune cells based on the disease status. **C,** Circle plots visualizing ligand-receptor interactions related to CCL pathway in sub-clusters of immune and epithelial cells using ‘CellChat’ package. **D,** Chord diagrams displaying significant interactions of the CCL pathway in epithelial cells with T_ex_ cell, MDSC, and macrophage sub-clusters. Directions from ligands (epithelial cell) to receptors (immune cell) are indicated. **E,** Tumors from Sox2-overexpressed PCN (PCNS) cell–transplanted mice were monitored and measured after intraperitoneal injection (days 17, 21, and 25) of Ccr2 inhibitors (CCR2 22 and BMS-813160) or DMSO (vehicle control); \*\*\**P* < 0.001, \*\*\*\**P* < 0.0001. **F,G,** Microscopic analyses of vehicle-, CCR2 22-, and BMS-813160-treated tumors; quantification (F) of IF images (G); scale bars = 20 μm, \**P* < 0.05, \*\**P* < 0.01, \*\*\**P* < 0.001, \*\*\*\**P* < 0.0001. **H,** Venn diagram showing overlapped transcription factors of the UCSC ChIP seq database and PN- and PCN-specific regulons. **I,** Rela regulon projection in regulon-based UMAP of WT, PC, and PCN datasets. The color shows regulon specificity score of the cells. **J,** Network analysis of transcription factors related to human Ccl2 gene expression. Target genes of transcription factors were refined by iRegulon, and the Ccl2 connections to Taf1 and Rela are displayed. **K,** Rela immunofluorescence images of 2D-cultured PN and PCN. Nuclei were stained with DAPI. Scale bar = 10 μm. The proportion of cells with nuclear-accumulated Rela was evaluated in PN and PCN cells. **L,** qRT-PCR results showing Ccl2 expression in PCN cells treated with different doses of NF-κB inhibitor for 6 hrs. **M,** ChIP assays showing binding activity of Rela to Ccl2 promoter in PN and PCN cells. Putative Rela binding sites (a, b, and c) and non-binding sites in the distant region (d) were analyzed with eluted DNA fragment amplification by PCR.

Given the role of the CCL2-CCR2 axis in the immune evasion of tumor cells,^30^ we tested whether the CCL2-CCR2 axis is indispensable for ESCC initiation using two different CCR2 inhibitors (CCR2 22 and BMS-813160). To increase tumorigenic efficiency, we established PCN cells that stably expressed Sox2 (PCNS), as Sox2 promotes ESCC tumorigenicity.^31^ Sox2 overexpression accelerated the tumor development of PCN cells while not affecting *Ccl2* expression (Figure S9A-C). Although CCR2 inhibitors did not suppress PCNS cell growth *in vitro*, allograft tumors grown with CCR2 inhibitors were drastically reduced in size and incidence compared to tumors treated with vehicle (Figure 5E, S9D). All tumors exposed to vehicle and CCR2 inhibitors showed abundant immune cell infiltration into the tumors and comparable numbers of proliferating (Mki67^+^) tumor cells (Figure S9E,F). Consistent with the *Ccl2* KO tumor results, T_ex_ cell (Pdcd1^+^) infiltration into CCR2-inhibited tumors was markedly decreased, while active CD8^+^ T cells (perforin^+^) were enriched in the tumors from CCR2 inhibitor–injected mice compared to in the tumors from vehicle-injected control mice (Figure 5F,G, S9F,G). On the other hand, fewer M2 macrophages were observed in the CCR2-inhibited tumors than in the *Ccl2* KO tumors (Figure 5F,G, S9F). However, the number of other monocyte-derived cells (CD68^+^ or CD209^+^) was lower in the CCR2 inhibitor–treated TME, indicating reduced monocyte recruitment by CCR2 blockade (Figure S9F-H). These results suggest that the CCL2-CCR2 axis elicits immune evasion for ESCC initiation.

### NF-**κ**B pathway, the upstream regulator of CCL2

Next, we determined the mechanism of CCL2 upregulation during ESCC tumorigenesis. Because CCL2 is upregulated mainly by transcription factors,^32^ we sought to identify the key transcription factors that transactivate *CCL2*. An scRNA-seq–based regulon analysis identified PCN-specific regulons (RELA, FOXA1, MYC, TAF1, and TCF12) that may bind to the *Ccl2* promoter (Figure 5H). Among them, Rela was the most highly activated in PCN, and *Ccl2* is expected to be directly transactivated by Rela from a refined analysis using iRegulon (Figure 5I,J). Compared to PN, PCN cells showed more abundant nuclear accumulation of Rela and Rela inhibition downregulated *Ccl2* in PCN cells (Figure 5K,L). Moreover, a chromatin immunoprecipitation (ChIP) assay showed that Rela occupied the *Ccl2* promoter in PCN but not in PN cells, consistent with the results of a previous report that RELA transactivates *CCL2* (Figure 5M).^32, 33^ In addition, a human ESCC tissue microarray analysis showed a positive correlation between CCL2 and RELA expression, supporting the pathological relevance of the RELA-CCL2-CCR2 axis to human ESCC tumorigenesis (Figure S9I,J).

### Classification of ESCC patients and PCN relevance

To extend our findings to the biology of human ESCC tumorigenesis, we analyzed single-cell transcriptomes of 69 ESCC patient tumor samples (Figure 6A) by integrating human scRNA-seq datasets with the PCN’s.^34, 35^ The unsupervised principal components analysis and Pearson correlation coefficient categorized 69 patient datasets into three groups (ESC1-3). While ESC1, 2, and 3 constituted a comparable portion of patients (ESC1: 44.9%; ESC2: 29.0%; ESC3: 26.1%) with tumor cell heterogeneity, as described previously,^34^ PCN belonged to ESC1 (Figure 6B,C). Notably, ESC1 displayed the lowest pathway score for TP53, CDKN2A, and NOTCH signaling, suggesting that ESC1 has consistent transcriptomic features of PCN EOs (Figure 6B, D). ESC1 and ESC2 showed the highest score for the NF-κB pathway, correlated with *CCL2* expression (Figure 6E,F). Consistent with PCN tumors’ immune landscape, tumor cells of ESC1 patients highly expressed *CD274*/*PD-L1*, suggesting that PCN-type (ESC1) likely has immune escape potential through PD-L1-PD1– mediated T cell exhaustion (Figure 6F, S10A). On the other hand, ESC2 exhibited activation of WNT, PI3K, MAPK, and FGFR1 pathways, whereas ESC3 was enriched with genes related to Hippo, cell cycle, and chromatin modification pathways (Figure S10B,C). Of note, despite the transcriptomic differences among ESC1-3, we could not find categorical differences between the conventional tumor stage classification in the ESC1-3 subtypes (Figure S10D).

**Figure 6.**
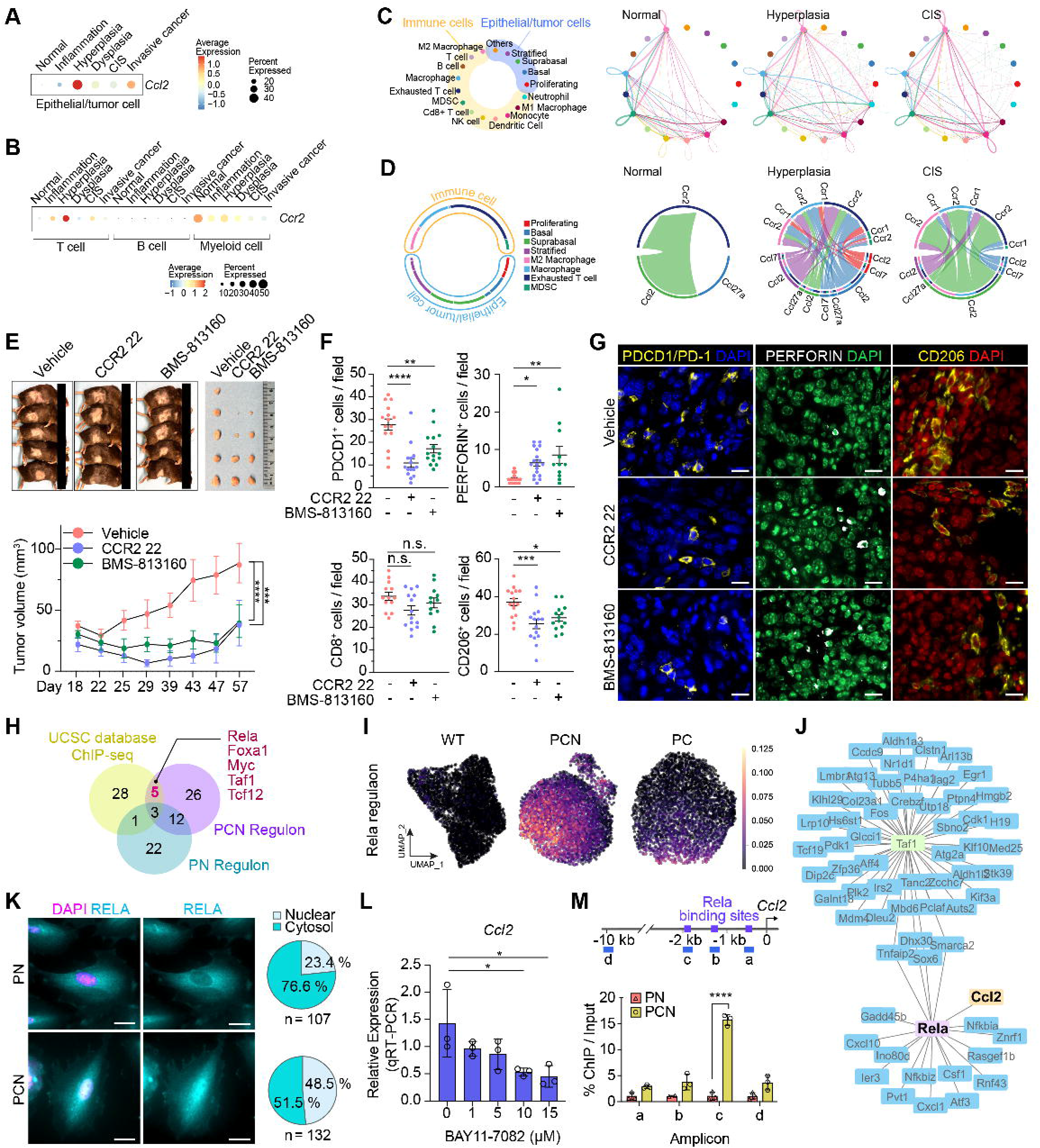
Classification of ESCC patients and PCN relevance. **A,** Integrated UMAP of tumor epithelial cells of 69 ESCC patients and PCN organoids. **B,** Correlation matrix heatmap of integrated datasets of ESCC patients and PCN organoids. The dendrogram showing the distance of each dataset based on principal component analysis with the Pearson correlation (a color spectrum) and pathway scores (TP53, CDKN2A, and NOTCH). PCN was excluded for pathway scores. **C,** UMAP showing ESCC subtypes. **D-F,** Dot plots of pathway scores (TP53, CDKN2A, and NOTCH) (D), NF-κB score (E), and *CCL2*, *CD274*/*PD-L1*, and *PDCD1LG2*/*PD-L2* gene expression (F) in ESCC subtypes. **G,** Dot plot of ESCC subtype’s representative marker genes selected from DEG analysis using the Wilcoxon method. **H,I,** B2M and CCL2 staining of human ESCC tissue microarray samples (**H**) and the correlation of the results (I); scale bars = 50 μm (upper) and 20 μm (lower); r: Pearson correlation coefficient; *P*: *P* values; N: number of samples. **J,K,** Heatmap of B2M staining results in tumor samples (J) and B2M, CCL2, and RELA staining results in ESCC patients with adjacent normal and tumor paired samples (K). Immunohistochemistry scores displayed from 1 (lowest) to 3 (highest) expression.

We further identified the representative marker genes of each group of patients by DEG analysis. ESC1 showed higher expression of *B2M*, *RGS1*, and *THEMIS2*, which were reported to be highly expressed in several types of cancer.^36–38^ *EID1*, *CYB5R3*, and *MGST3* were highly expressed in ESC2, and *MFL2*, *FUS*, and *MDK* were enriched in ESC3 patients (Figure 6G, S10E). Although these markers are upregulated in various cancers,^39–41^ they have not been investigated in ESCC. ESCC patients with higher expression of *B2M*, *RGS1*, and *FTL* exhibited poor survival than did those with lower expression (Figure S10F). A TCGA-based analysis also showed that *B2M*, *RGS1*, *MLF2*, *FUS*, and *MDK* were upregulated in ESCC compared to in normal samples (Figure S10G). Immunostaining of the ESCC tissue microarray (*n* = 101) revealed a positive correlation between B2M and CCL2 expression and B2M and RELA expression (Figure 6H,I, S10H). Most patients (90.9%) had relatively higher B2M expression in tumor samples than in adjacent normal samples, and 22.7% (5 of 22, immunohistochemistry score ≥ 2.5) showed robust expression of both CCL2 and B2M in the tumor tissue (Figure 6J). Focusing on tumor samples alone, 49.0% (25 of 51) of tumor samples expressed high levels of B2M (immunohistochemistry score ≥ 2.5), a similar proportion to ESC1 from scRNA-seq (Figure 6K). These data suggest that B2M is a prognostic marker of ESCC, likely with RELA-CCL2 axis activation.

## Discussion

To identify the multiple genetic factors responsible for ESCC initiation, we utilized organoids, which overcome the limitation of single-gene KO (e.g., CRISPR-based screening) and provide a better phenotype analysis (i.e., neoplasia), as further confirmed by a congenic transplantation approach. Inactivation of *Trp53*, *Cdkn2a*, and *Notch1* (PCN) was sufficient to initiate ESCC. While PN and PCN EOs displayed the neoplastic features of ESCC *in vitro*, only PCN cells exhibited *in vivo* tumorigenicity, suggesting immune evasion driven by PCN. A transcriptomic analysis identified that the CCL2-CCR2 axis remodels the immune landscape, confirmed by loss-of-function studies (*Ccl2* KO and CCR2 inhibitors) and a scRNA-seq analysis of the autochthonous mouse model. Finally, using single-cell transcriptomics, we stratified ESCC patients and identified the subtype of PCN-type ESCC patients with new molecular signatures.

Tumor cells evade immune surveillance in various ways.^42, 43^ Despite several studies describing the immune landscape of ESCC,^34, 35, 44, 45^ experimental demonstrations of ESCC immune evasion have not been achieved. Our comprehensive approaches revealed that *CDKN2A* inactivation plays a crucial role in establishing tumor-favorable niches. Immune profiling of the mouse ESCC model showed decreased effector T cells and increased immune suppressive dendritic cells during ESCC development (Figure 4J,K, S7E,F).^29^ Similarly, human ESCC generates an immune-suppressive landscape with enriched T_ex_ or T_reg_ cells and upregulation of immunosuppression-related genes in TAMs and dendritic cells.^34, 35, 46^ Notably, enriched T_ex_ cells and TAMs are commonly observed in mouse scRNA-seq–based immune profiling and PCN TME (Figure 4J,K, S7E,F), unveiling the distinct immune niche of PCN-associated ESCC tumorigenesis.

We found that CCL2 is the key immune landscape remodeling factor for ESCC initiation, mainly represented by increased T_ex_ cells and M2-like macrophages and a reduced number of effector CD8^+^ T and dendritic cells. The CCL2-CCR2 axis induces tumor-favorable TME.^47^ Given the high expression of CCR2 in monocytes, CCR2-mediated M2-like macrophage differentiation and polarization has been studied in the context of tumor cell invasion and metastasis.^27^ Additionally, the CCL2-CCR2 axis was shown to induce TAM accumulation.^30^ Moreover, CCR2 expressed in T cells modulates the recruitment of T_reg_ cells,^48^ hampering immune surveillance. Therefore, targeting the CCL2-CCR2 axis to enhance immune surveillance may be a promising option for ESCC treatment. Indeed, blocking the CCL2-CCR2 axis has been tested in clinical trials in other types of cancer, with ICIs.^28, 47, 49^ However, since CCL2 is a chemoattractant that recruits monocytes, systemic inhibition of the CCL2-CCR2 axis might affect normal immune surveillance. For instance, impaired monocyte recruitment by CCR2 blockade could reduce the number of antigen-presenting cells, diminishing innate and adaptive immune responses critical for anti-cancer effects.^50^ Our results showed that the transient inhibition of CCR2 suppressed tumor growth, but monocyte-derived cell (CD209^+^ or CD68^+^) recruitment was decreased. In contrast, the number of CD8^+^ T cells was not significantly increased compared to the TME of *Ccl2* KO tumors (Figure 5, S9). Therefore, targeting CCL2 or the upstream molecules of CCL2 in tumor cells would minimize potential adverse effects.

Recent clinical trials showed promising outcomes of ICIs in patients with advanced ESCC.^10–14^ Nonetheless, ESCC patient stratification is a significant hurdle. Although the datasets from the 69 patients analyzed in this study lacked ICI-related clinical information, PCN-type (ESC1) patients showed higher expression of *PD-L1*/*CD274* than did other patients. Thus, CCL2-CCR2 blockade combined with PD-L1 or PD-1 ICIs could be effective in ESC1 patients, and B2M could be a signature for ESC1 subtype (Figure 6G, S10E), potentially allowing a molecular signature-guided ICI strategy.

Together, this study identifies the inactivation of *TP53*, *CDKN2A*, and *NOTCH1* as important genetic events leading to ESCC development, accompanied by immune evasion, and newly classifies ESCC patients based on the molecular features of tumor cell transcriptomes.

## Data and code availability

Bulk and scRNA-seq data are available via the Gene Expression Omnibus (GEO; GSE213929; log-in token for reviewers: yzcpewsqddwpzcf). The code can be accessed via GitHub (https://github.com/jaeilparklab/ESCC_project_1) and is available upon request.

## Supporting information

Supplementary Table 1. Primer information

Supplementary Table 2. single guide RNA_sequences of genes

Supplementary Table 3. Single-cell datasets information

Supplementary Table 4. Patients ID and pathologic stages

Supplementary Table 5. Cell numbers of each cell type

Supplementary Table 6. DEG from RNA-seq

Supplementary Table 7. DEG from scRNA-seq

Supplementary Table 8. Gene lists for pathway scores_v02

## Acknowledgments

We are grateful to Kwon-Sik Park, Pierre D. McCrea, and Malgorzata Kloc for comments; Ann Sutton (Research Medical Library, MD Anderson) for editing the manuscript; and the Herbert Irving Comprehensive Cancer Center for the shared resources (Biostatistics, Genomics, and Molecular Pathology, and 3D organoid/cell culture). This work was supported by the Cancer Prevention and Research Institute of Texas (RP200315 to J.-I.P.), the National Cancer Institute (CA193297 and CA256207 to J.-I.P; 5P30 CA013696 and 5P01 CA098101 to A.-K.R., H.N., K.D., G.E., C.M.), an Institutional Research Grant (MD Anderson to J.-I.P.), a Specialized Program of Research Excellence (SPORE) grant in endometrial cancer (P50 CA83639), and Radiation Oncology Research Initiatives. The core facilities at Baylor College of Medicine (Cytometry & Cell Sorting Core and Single Cell Genomics Core) were supported by CPRIT (RP180672, RP200504) and the National Institutes of Health (CA125123, RR024574).

---

Author names in bold designate shared co-first authorship.

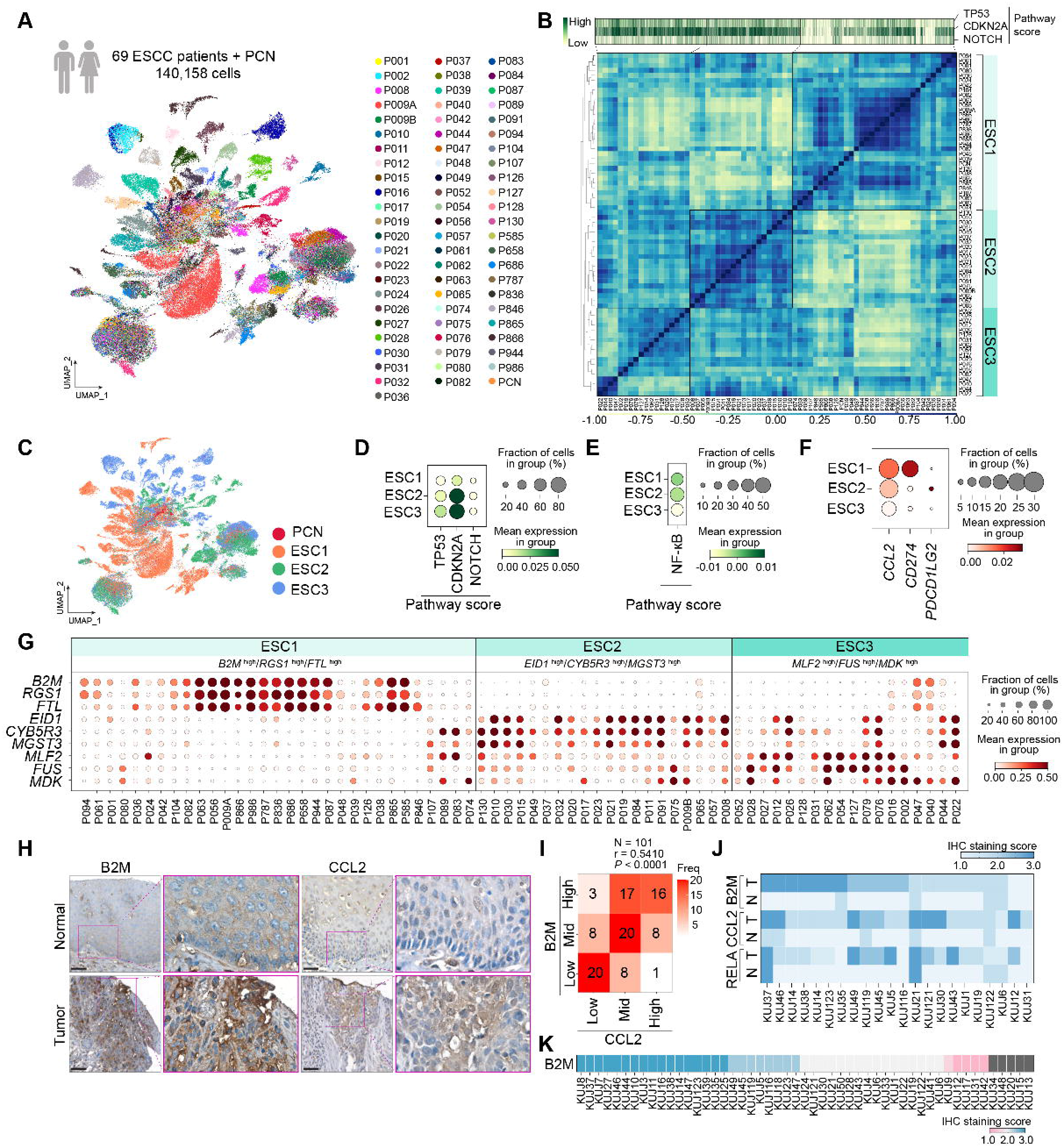

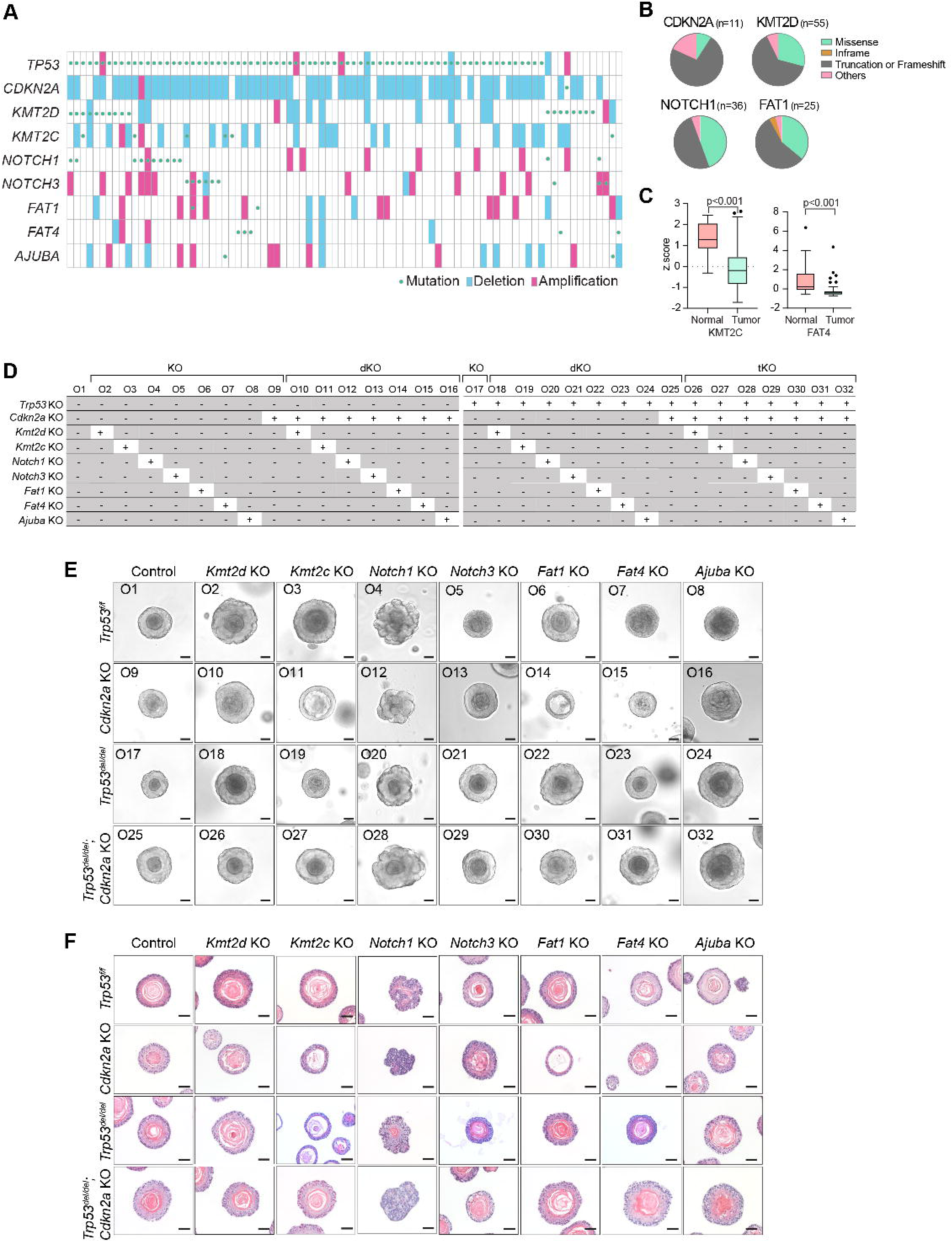

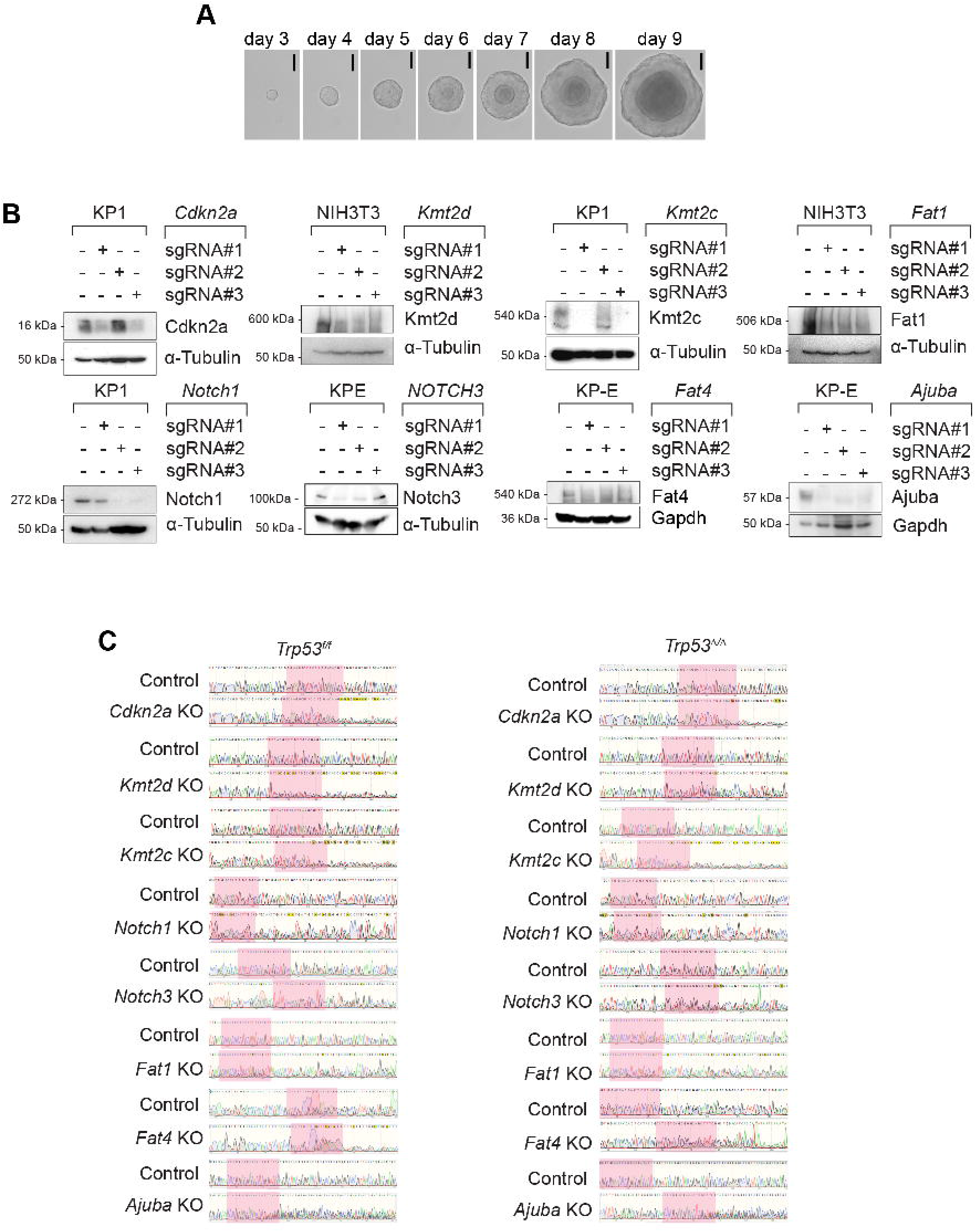

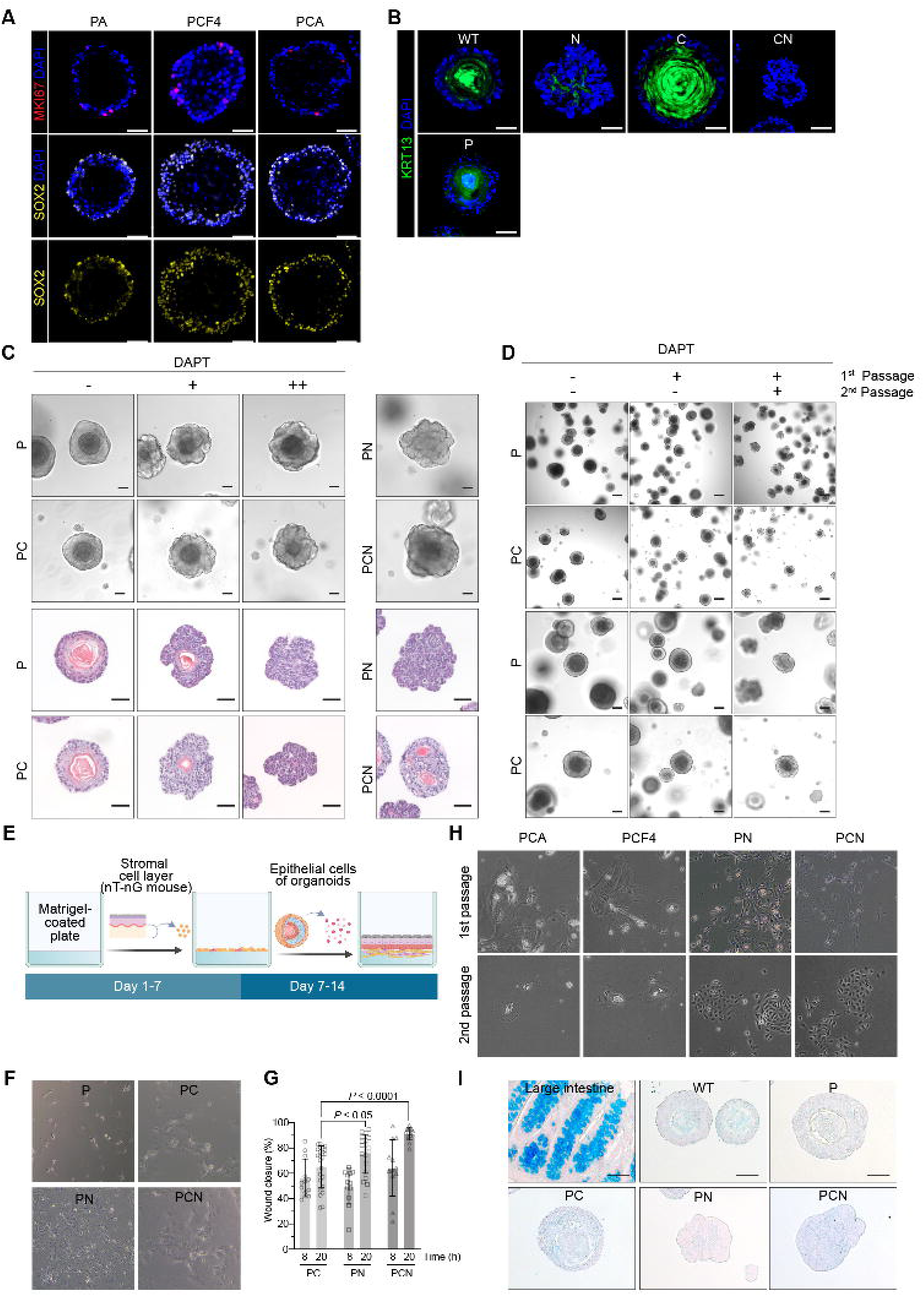

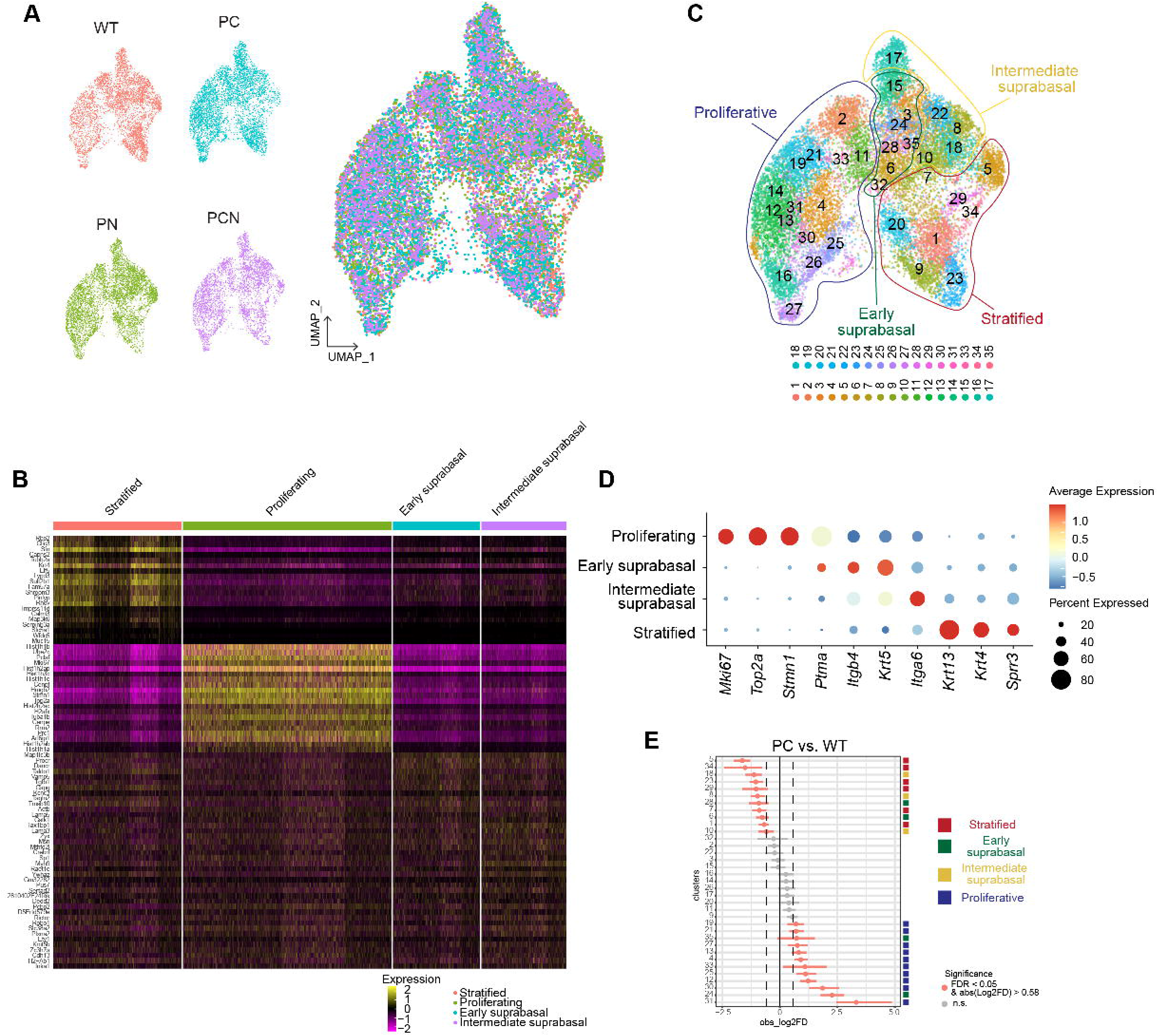

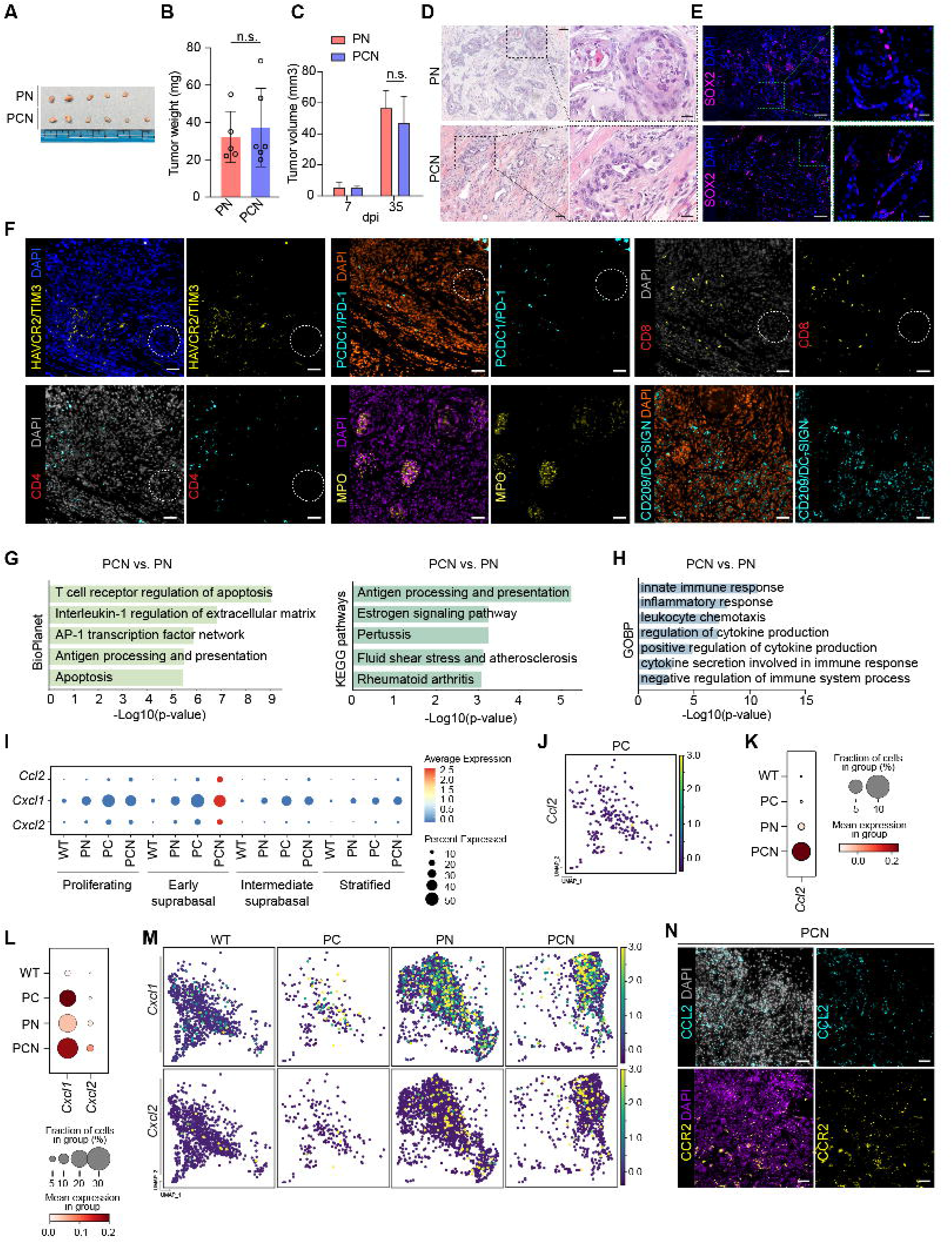

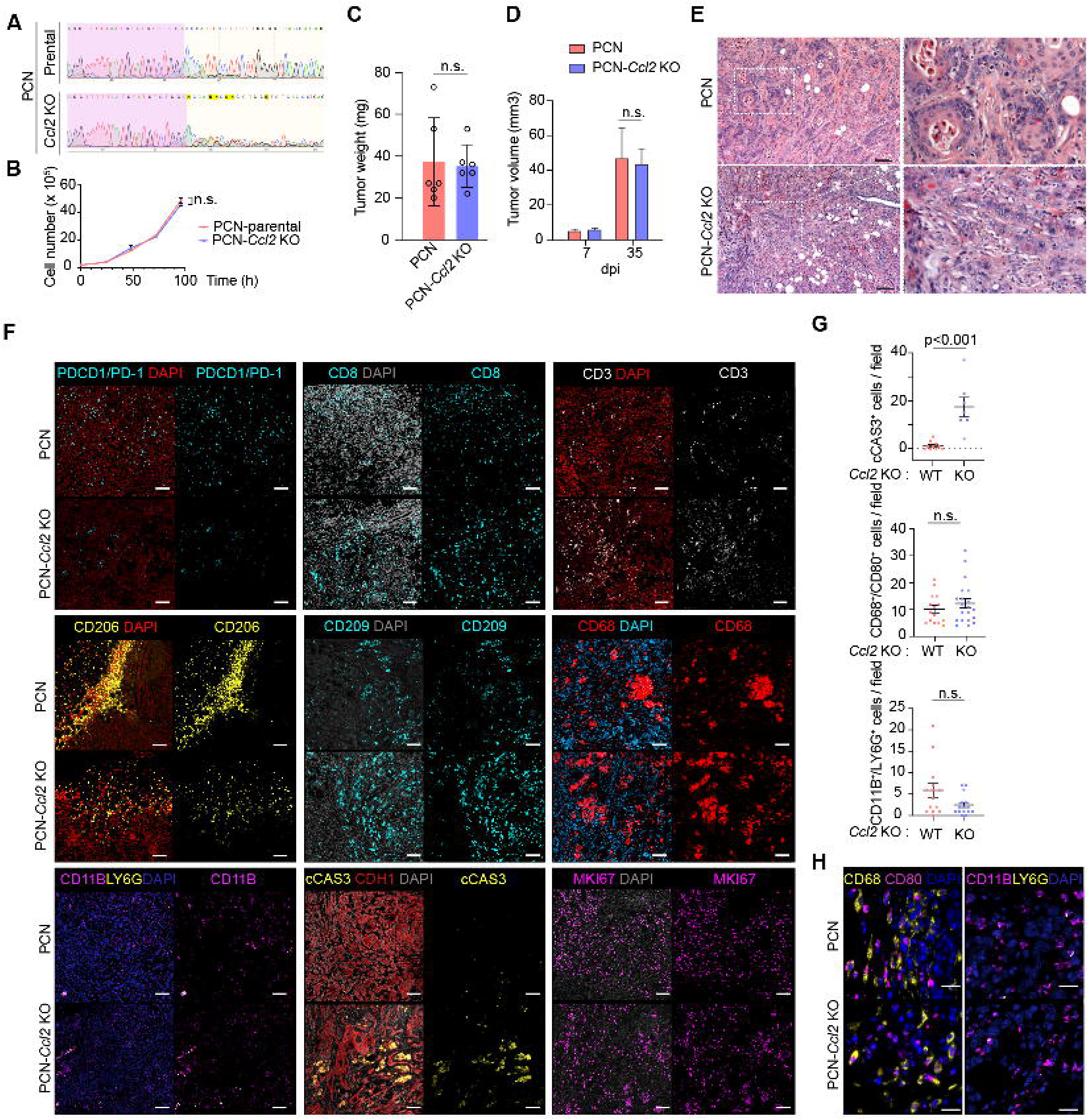

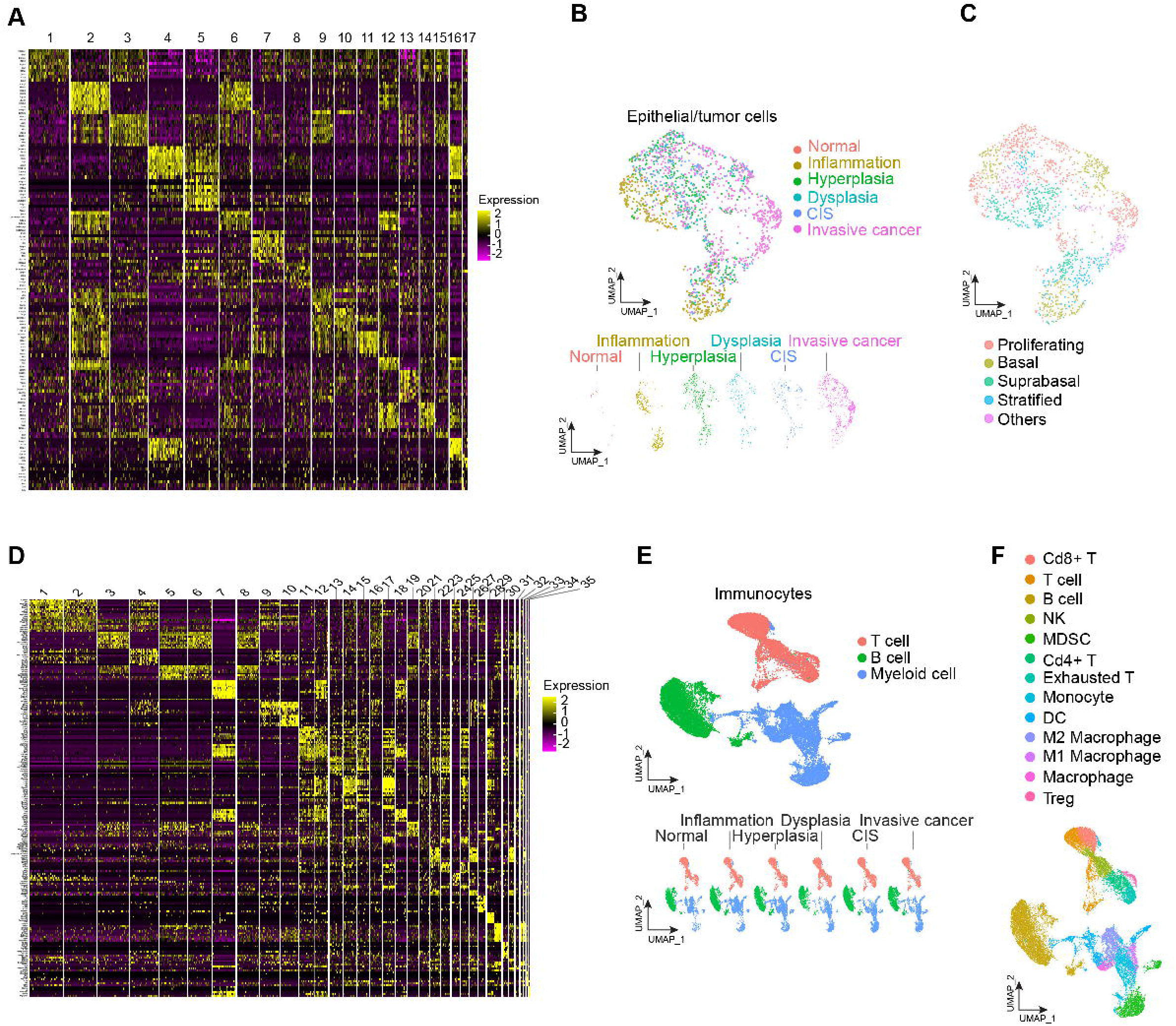

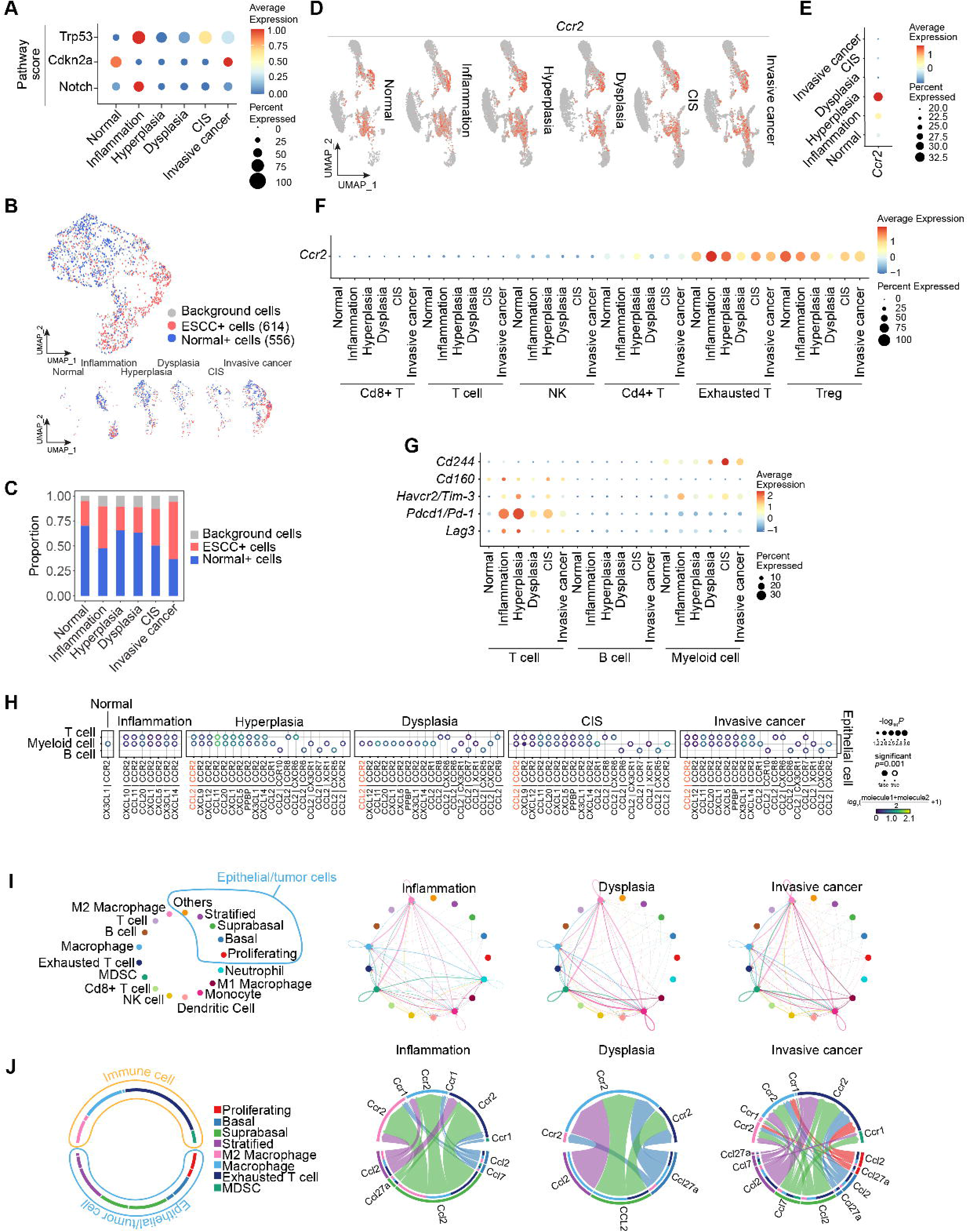

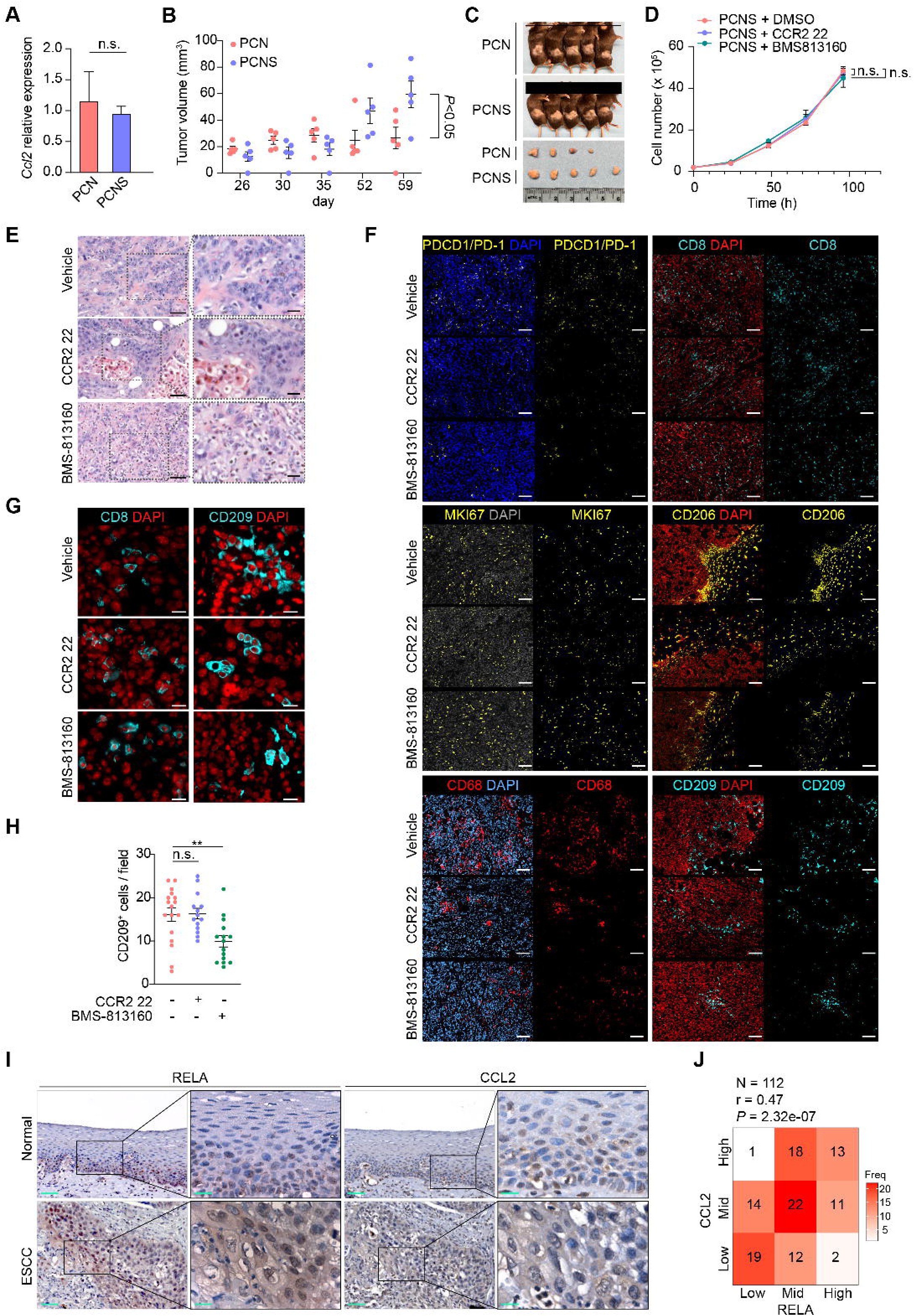

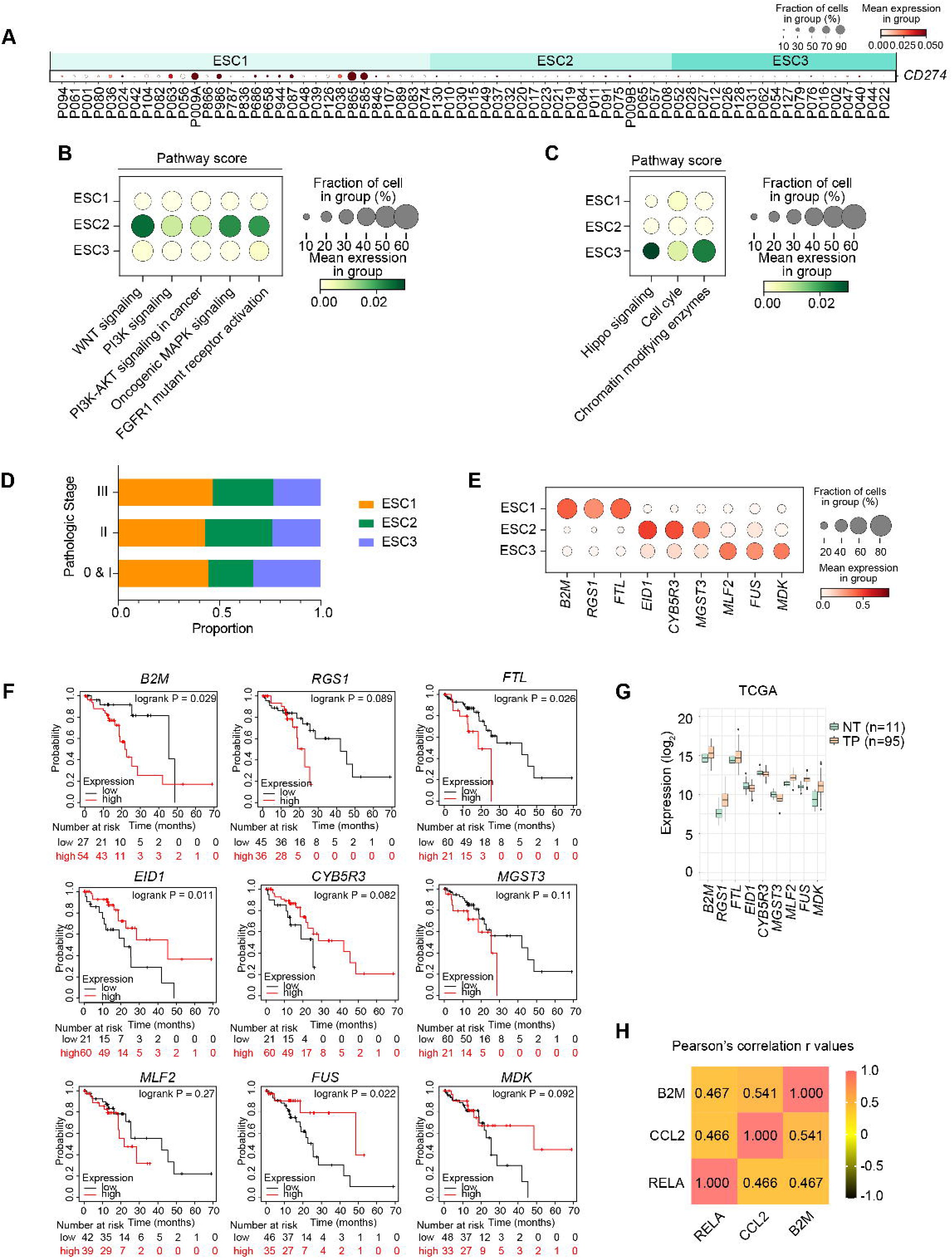

