## Supplementary Methods for "Key Genetic Determinants Driving Esophageal Squamous Cell Carcinoma Initiation and Immune Evasion"

**Wnt3A, R-Spondin1, and Noggin-conditioned medium**

The Wnt3A, R-Spondin1, and Noggin (WRN)-conditioned medium was prepared as previously described.^1^ In brief, L­WRN (ATCC CRL­3276) cells were cultured on a 10-cm plate with culture medium (Dulbecco’s modified Eagle’s medium [DMEM, Fisher], 0.5 mg/mL G418 (Thermofisher), 0.5 mg/mL hygromycin B (Thermofisher), 1% penicillin/streptomycin (Life technologies), and 10% fetal bovine serum [FBS]). After 10% of L-WRL cells (ATCC, CRL-3276) had been seeded in culture medium (without G418 and hygromycin B) in 10-cm plates, cells were incubated for 3-4 days. The medium was replaced with 10 mL of fresh medium when the cells were 80%-90% confluent, and the cells were incubated for 24 hrs. The medium was collected, centrifuged at 1000 × g for 4 min, passed through a 0.22-μm sterile filter, and stored at −80 °C. Another 10 mL of fresh medium was added to the plates and collected after 24 hrs to make the second batch of conditioned medium using the same procedures. The first, second, and third batches of conditioned media were mixed before use to prepare a 100% WRN-conditioned medium.

**Esophageal tissue isolation**

The cervical dislocation was performed after the 8- to 10-week-old mice had been euthanized via CO2 inhalation. The esophagi were collected in 10-cm Petri dishes with ice-cold phosphate-buffered saline (PBS) with 1% penicillin/streptomycin and swirled gently to remove blood. The esophagi were opened longitudinally and washed with cold PBS with 1% penicillin/streptomycin. The epithelial cell layer was peeled off using surgical tweezers and then dissected into 0.5-cm3 pieces with surgical blades. The minced esophagi were collected in a 15-mL conical tube and digested by 0.05% trypsin-EDTA (Thermofisher) at 37 ℃ for 60 min with frequent vortexing. After dissociation, 3× volume 10% FBS-supplemented DMEM was added to inactivate the dissociation enzyme, followed by vigorous pipetting. The suspension was passed through a 35-μm sterile cell strainer to collect a single-cell suspension. Finally, the cell suspension was spun down at 1000 rpm for 4 min at ambient temperature and resuspended in a 50% WRN-conditioned medium.

**EO culture**

The E-MEOM included 50% WRN medium (50% advanced DMEM/F12 [Thermofisher], 50% WRN-conditioned medium, 1% penicillin/streptomycin, and 1 × GlutaMAX [Life Technologies]), 1 × B27 (Thermofisher), 50 ng/mL, 10 mM [nicotinamide](https://www.sciencedirect.com/topics/biochemistry-genetics-and-molecular-biology/nicotinamide) (Sigma), 500 nM A83-01 (Sigma), 10 μM SB202190 (Sigma), 50 ng/mL EGF (PeproTech), and 10 μM Y-27632 (Thermofisher) ROCK inhibitor (first 3 days). Single-cell dissociated esophageal epithelial cells (1000 cells) were suspended in 8 μL of E-MEOM and 12 μL of pre-thawed Matrigel (Corning) on ice and seeded in the centers of a well to create a Matrigel dome. The plate was incubated at 37 °C for 10 min to solidify the Matrigel. Finally, 500 μL of E-MEOM was added and incubated at 37 °C with 5% CO2. The medium was changed every 2-3 days.

***Trp53 KO* EO**

8- to 10-week-old *Trp53^floxed/floxed^* mice were euthanized to collect the esophagi, which were digested into a single-cell suspension and seeded in Matrigel to form EOs, as described above. After 7 days of culture, EOs were digested with 0.05% trypsin-EDTA at 37 °C for 45 min to create a single-cell suspension. Ad-CMV-EGFP or Ad-Cre-EGFP (University of Iowa) was added to the cell suspension at 1 × 103 pfu/cell. Cells were then suspended in Matrigel to generate new EOs. After 2 days of seeding, 10 μM nutlin3 was added for *Trp53^floxed/floxed^* cell selection. Two days later, the selection was performed by sorting the GFP+ cells after dissociating EOs into single cells, and the sorted cells were re-seeded for EO culture as described above. EOs were collected for genotyping 7 days after seeding. Wild-type, *Trp53^floxed/floxed^*, and KO Trp53 alleles were amplified as 288 bp, 370 bp, and 612 bp, respectively. See Supplementary Table 1 for primer information.

**CRISPR/Cas9-based gene KO in EOs**

CRISPR/Cas9 system-mediated gene KO was described in a previous study.^2^ In brief, WT or Trp53 KO EOs were digested with 0.05% Trypsin-EDTA to dissociate into single cells. Single cells were incubated with a virus-containing medium with polybrene for 1 hr with centrifugation (600 g) at 32 °C (see Supplementary Table 2 for sgRNA sequences). Cells were then incubated at 37 °C with 5% CO2 for 4 more hrs and embedded in Matrigel. The medium was replaced 2 days after infection with antibiotics (puromycin [Sigma], blasticidin [Invitrogen], or hygromycin) for selection. Gene KO was confirmed by genomic DNA PCR after lysis of the organoid using specific primer pairs (see Supplementary Table 1 for primer information).

**Sox2 stable cell line establishment**

Sox2-overexpressing PCN cells were established by lenti-viral transduction. Briefly, pLenti-Sox2-GFP plasmid was prepared by amplifying the *Sox2* template from the pLV-tetO-Sox2 plasmid (addgene, #19765), and then transferred to pLenti CMV GFP plasmid (addgene, #17446) using High-Fidelity DNA polymerase (NEB). Viral transduction was performed as described in a previous study.^2^ GFP-expressing cells were isolated using flow cytometry (BD FACSAria II Cell Sorter) and cultured again.

**Organoid-forming efficiency and size analysis**

After 7 days of organoid seeding in Matrigel, the size of the organoids was analyzed by measuring the volume under the microscope (ZEN software, ZEISS). To reduce the vulnerability of EOs, the measurements were performed after at least 3 passages after isolation from the KO experiments. All experiments included more than 50 organoids per group.

**H&E, PAS, and immunofluorescence staining**

All staining was performed as previously described.^3^ 7 days after seeding, EOs were collected by dissociating Matrigel using ice-cold PBS and fixed in 4% paraformaldehyde at ambient temperature. For tumor tissue, excised tumors were washed with ice-cold PBS and fixed with formaldehyde at ambient temperature. After paraffin embedding, tumor tissue and organoid sections were mounted on glass slides. For H&E staining, sections were incubated in hematoxylin for 3-5 min and eosin Y for 20-40 s. For PAS staining, slides were immersed in the periodic acid solution (Sigma) for 5 min at ambient temperature and then in Schiff’s reagent (Sigma) for 15 min at ambient temperature, followed by hematoxylin solution for 60-90 s. After washing with tap water, slides were dried, and the coverslips were mounted with mounting media. For immunofluorescence, after blocking with 3% goat serum in PBS for 30 min at ambient temperature, sections were incubated with primary antibodies (KRT13 [Abcam, ab92551, 1:250], MKI67 [Abcam, ab16667, 1:250], SOX2 [Cell Signaling Technology, #3579, 1:250], TRP63 [Abcam, ab124762, 1:250], cleaved Caspase-3 [Cell Signaling Technology, #9664, 1:250], RELA [Cell Signaling Technology, #8242, 1:200], PDCD-1/PD-1 [Cell Signaling Technology, #84651, 1:200], PERFORIN [Cell Signaling Technology, #44865, 1:200], CD8 [Cell Signaling Technology, #98941, 1:200], CD206 [Cell Signaling Technology, #24595, 1:200], CD209 [Thermofisher, PA5-119030, 1:200], HAVCR2/TIM3 [Cell Signaling Technology, #83882, 1:200], CD4 [Cell Signaling Technology #25229, 1:200], MPO [Thermofisher, #PA5-16672, 1:200], CCL2 [R&D Systems, MAB479-100, 1:200], CCR2 [R&D Systems, MAB55382, 1:200], CD3 [Cell Signaling Technology, #99940, 1:200], CD68 [Cell Signaling Technology, #97778, 1:200], CD11B [Cell Signaling Technology, #17800, 1:100], LY6G [Cell Signaling Technology, #88876, 1:100], and CD80 [Proteintech, 66406-1-Ig, 1:200]) overnight at 4 °C and secondary antibody (1:250) for 1 hr at ambient temperature. Sections were mounted with ProLong Gold antifade reagent with DAPI (Invitrogen). Images were captured with the fluorescence microscope (ZEISS; AxioVision and ZEN software).

**Immunohistochemistry**

ESCC cancer tissue microarray slides contained 144 samples from 55 patients (provided by Dr. Hiroshi Nakagawa). Immunostaining was performed as previously described ^3^. In brief, paraffin-embedded tissue antigens were retrieved with a basic or citrate antigen retrieval buffer. After being blocked with goat serum in PBS, tissues were incubated with primary antibodies (CCL2/MCP-1 [Thermofisher, MA5-17040, 1:200], RELA, and B2M [Cell Signaling Technology, #12851, 1:200]). The immunohistochemistry results were scored, analyzed, and visualized using the “gglot2” package in r and GraphPad Prism (v9.2.0).

**Gene expression analysis**

Organoids were harvested and lysed, and RNAs were extracted by TRIzol reagent. RNA was quantified with a Nanodrop 2000c Spectrophotometer (Thermo Scientific) and then converted to cDNA using SuperScript II reverse transcriptase with random hexamers. PCR amplification (StepOne Real-Time PCR System, Applied Biosystems) was performed using the following conditions: 95 °C for 10 min, 95 °C for 15 s (denature), and 60 °C for 1 min (anneal/extend) for 40 cycles; 95 °C for 15 s and 60 °C for 1 min; and 95 °C for 15 s (melting curve). qRT–PCR results were quantified by comparative 2^−ΔΔCt^ methods (see Supplementary Table 1 for primer information). The results were expressed as average fold change in gene expression and were normalized to the expression of *mHprt*.

**BrdU incorporation assay**

Organoids were incubated with BrdU (Cayman, 10 μM) for 0.5 hr from 7-day cultured organoids. BrdU-containing medium was removed, and Matrigel-embedded organoids were harvested. EOs were washed three times with PBS and fixed with 4% PFA for 30 min at ambient temperature. After the paraffin-embedding process, slides were prepared for staining. Following the immunofluorescence staining method, the slides were incubated with anti-BrdU antibody (Abcam, ab6326, 1:200) at 4 °C and fluorescence-conjugated secondary antibody. Samples were further stained with DAPI and mounted for imaging.

**Organotypic co-culture**

Four-well chamber slides (Falcon) were coated with 200 μg/mL Matrigel and stored at 4 °C until cell seeding. Stromal cells from esophagi were isolated from Rosa26nT-nG mice. We collected the stromal cell layer by peeling off the epithelial cells from the longitudinally cut esophagi. Stromal cells were dissociated with collagenase/dispase (Sigma) for 1 hr at 37 °C and passed through a 70-μm cell strainer. After counting, cells (1.2 × 10^5^ cells/well) were seeded in the pre-warmed Matrigel-coated chamber. Cells were grown with DMEM supplemented with 20% FBS for 14 days. Epithelial cells were collected from WT, PN, and PCN organoids and seeded on the stromal cell–cultured chamber slide. The medium was replaced every 2 days and cultured for 14 days. For staining, cells were fixed with 4% PFA for 30 min at ambient temperature and then incubated with anti-CDH1 antibody (Cell Signaling Technology, #3195, 1:200) overnight at 4 °C. Cells were visualized using secondary antibodies conjugated with fluorescein (1:200) for 1 hr at ambient temperature, and the images were taken with a fluorescence microscope (ZEISS; AxioVision and ZEN software).

**2D culture**

EOs were digested with 0.05% trypsin-EDTA at 37 °C for 45 min to create a single-cell suspension in a 15-mL centrifuge tube. The cells were centrifuged down at 1000 rpm for 4 min at ambient temperature, suspended in DMEM + 10% FBS with 10 μM Y-27632, and seeded on a 24-well plate. Cells were passaged every 3-5 days. After the third passage, Y-27632 was removed from the culture medium. DMEM supplemented with 10% FBS and 10% DMSO was used to freeze cells and store them in liquid nitrogen.

**Colony formation, cell growth, wound-healing assays**

Cells (1 × 10^4^) were seeded on a 60-mm dish, and the medium was replaced every 2 days. The cells’ colony-forming ability was monitored, and they were fixed with methanol for 20 min. The fixative was removed, and the dishes were rinsed with distilled water. The colonies were stained with 0.05% crystal violet solution, and the dishes were dried after being washed three times with distilled water. For the cell growth analysis, 2 × 10^5^ cells were seeded on the 60-mm dish. The cell number was measured daily using an automated cell counter (Biorad) after the cells were trypsinized and stained with Trypan Blue. For the wound-healing assay, 1 × 10^4^ cells were seeded on the 24-well plate. A wound was made in the middle of the well after the cells were confluent, and the cells were imaged at 0, 8, and 20 hr.

**Xenograft and allograft transplantation**

Five-week-old C57BL/6 scid mice were purchased from the Jackson Laboratory (#001913) and C57BL/6 mice were maintained in the Division of Laboratory Animal Resources facility at MD Anderson. 2D-cultured PN, PCN, PCN-*Ccl2* KO, PCNS, and PCNS-*Ccl2* KO cells (5 x 10^5^ for scid mice and 3 × 10^6^ for C57BL/6 mice) were injected subcutaneously into the right dorsal flanks of mice, respectively. Tumor volume was calculated by measuring with calipers every 3-4 days (volume = (length × width2)/2). For CCR2 inhibitor treatment, DMSO-dissolved BMS CCR2 22 (1 mg/kg) and BMS-813160 (25 mg/kg) were injected intraperitoneally at day 17, 21, and 25. DMSO was injected as vehicle control. Mice were euthanized, and tumors were collected at day 57. The excised tumors were photographed and paraffin-embedded for immunostaining.

**Library preparation for RNA-seq and scRNA-seq**

For bulk RNA-seq, triplicated WT, PN, and PCN organoids were harvested 7 days after seeding and lysed using the RNeasy Plus Mini Kit (QIAGEN). Purified RNA was used for mRNA library preparation, sequenced by Illumina NovaSeq, and mapped to GRCm38/mm10 genome (Novogene).

For scRNA-seq, organoids from WT, PC, PN, and PCN were collected 7 days after seeding and digested with 0.05% trypsin-EDTA at 37 °C for 30 min. After trypsin had been inactivated with 10% FBS DMEM, a single-cell suspension was collected by passing cells through a 35-μm cell strainer. Each group was tagged with two CMO tags from the CellPlex kit (10× Genomics). The tagged cells of each group were pooled together with the same number of cells after being counted. The cDNA library was prepared with the 10× Genomics 3’ v2 kit and sequenced on an Illumina NovaSeq (Novogene), mapped to the GRCm38/mm10 genome, and demultiplexed using CellRanger (v7.0.1). The resulting count matrices files were analyzed in R (Seurat [v4.0.3]) or Python (Scanpy [v1.8.2]).

**Public scRNA-seq data preparation**

General information of scRNA-seq datasets was described in Supplementary Table 3.

**Mouse datasets.** Mouse 4-NQO-treated scRNA-seq datasets were obtained from the Genome Sequence Archive in BIG Data Center (Beijing Institute of Genomics, Chinese Academy of Sciences, http://gsa.big.ac.cn) under the accession number CRA002118. Mouse datasets were directly used for analysis as downloaded, with filtered matrices.

**Human datasets.** ESCC patient datasets were downloaded from the National Center for Biotechnology Information Sequence Read Archive under the accession numbers PRJNA777911 and PRJNA672851. ESCC patient datasets were downloaded and converted to fastq files using Sequence Read Archive tools and the parallel-fastq-dump package. The converted fastq files were used as input for the CellRanger (v6.1.2) pipeline. Analyses were conducted with CellRanger output files. The datasets from 11 patients (PRJNA777911) and tumor samples that had been previously sorted by CD45-negative selection from 58 patients (PRJNA672851) were input to run CellRanger (v7.0.1). Patient ID and tumor grade are described in Supplementary Table 4.

**scRNA-seq data analysis**

**integration and clustering.** Organoid scRNA-seq data analysis. For pre-processing and clustering of scRNA-seq data, we used the Seurat R package and Python package Scanpy. For the organoid datasets, since two tags were used for one group of organoids, two datasets of the same genotype were integrated using “sctransform” in Seurat and “concatenate” in Scanpy and then annotated as WT, PC, PN, or PCN. The batch correction was performed in Scanpy using “Harmony”. UMAP was used for dimensional reduction, and cells were clustered into 35 groups in Seurat. Each cluster was annotated based on marker gene information. Seurat was used for 4-NQO-treated mice datasets that had been sorted with CD45^-^ and CD45^+^ cells. Datasets were pre-processed, normalized separately, and annotated based on their marker gene expression. Disease status was annotated by converting the original implemented identity class, “X0W, X12W, X20W, X22W, X24W, X26W” to “Normal, Inflammation, Hyperplasia, Dysplasia, CIS, and Invasive cancer”. Scanpy was used for human datasets preprocessing and integration. Each patient dataset was normalized separately and clustered by the “Leiden” algorithm. The clusters expressing a high level of PTPRC (CD45) were removed to exclude immune cell contamination and then a total of 69 patient datasets were integrated using the “concatenate” function. Batch effects were corrected using “Harmony”. See Supplementary Table 5 for cell numbers in each cell cluster.

**Proportion difference test.** The differences between the clusters from the two datasets were tested using the scProportionTest (v0.0.0.9) package. The cluster difference between the two datasets was compared, and the significance was calculated from the *P* value and confidence interval for the magnitude difference via bootstrapping using the default parameter of permutation (n = 1000).

**Trajectory inference.** RNA velocity was used for cell lineage tracing and latent time inference based on the previous report.^4^ Bam files produced from the CellRanger (v7.0.1) pipeline were used to create Loom files. Using the Python-based velocyto package, we produced Loom files that included unspliced and spliced read information. Two Loom files were generated for one genotype of organoid since the multiplex process had used two oligo tags for each genotype. The Loom files from the same genotype were merged using the “combine” function in the Loompy (v3.0.6) package. Cells were filtered, and dimensional reduction was performed following the default parameters using the scVelo (v0.2.4) and Scanpy packages. RNA velocity was calculated through a dynamical model, and cells were clustered using the “Louvain” algorithm. RNA velocity for all four datasets was performed with the same parameters (n_neighbors=10, n_pcs=50). Latent time analysis and a PAGA analysis were performed and plotted using the scVelo package.

**Enrichr analysis.** To perform Enrichr analysis of PCN- or PN-enriched genes, we performed DEG analysis from bulk RNA-seq and scRNA-seq results. DESeq2 (v1.30.1) package was used for bulk RNA-seq and PCN highly expressed 1516 genes were used as input gene list. For scRNA-seq data, ‘FindMarkers’ function was used to make the DEG list from proliferating cell clusters of PCN and PN datasets. 100 genes highly expressed in PCN compared to PN were used for Enrichr analysis. The analysis was conducted following instructions as described.^5^ See Supplementary Table 6 and 7 for DEG results.

**fGSEA analysis.** To perform a GSEA of PCN- or PN-enriched genes with human ESCC samples, we prepared a rank gene list from the DEG analysis of human ESCC vs. normal RNA-seq data (TCGA). The “glmLRT” method was used, and the fdr cut-off was 0.05. We collected 495 ESCC highly expressed genes and 258 normal highly expressed genes and filtered the not-assigned genes, resulting in 750 genes for analysis. The geneset was created from PCN or PN organoid highly expressed genes compared to WT organoid-expressed genes from scRNA-seq data. To match the names of genes in the rank gene list and geneset, we converted mouse gene names to human gene names using the biomaRt (v2.46.3) package. In total, 2812 PCN and 1285 PN highly expressed genes were used to create genesets. The enrichment value was calculated and plotted with the fgsea (v1.16.0) package (permutation number = 2000).

**Similarity test.** The Scissor package was used to compare the transcriptome of the organoids scRNA-seq dataset and the bulk RNA-seq datasets of ESCC patients.^6^ To compare scRNA datasets with ESCC patients, we downloaded ESCC patient data and normal sample HTSeq count data from the GDC data portal (TCGA-ESCA). After preparing the normal and ESCC gene expression matrix, we converted the human gene names to mice gene names using the biomaRt (v2.46.3) package. Not converted or assigned genes and duplicated gene names were removed from the matrix, and a Scissor analysis was performed with each scRNA dataset using the Cox regression model (alpha = 0.01). To compare scRNA-seq data with poor survival patient data, we downloaded ESCC patients’ clinical metadata from the GDC data portal and binarized it into “poor survival” and “better survival”. A Scissor analysis was conducted with scRNA datasets using the Cox regression model and 0.03 alpha value.

**Regulon analysis.** For the regulon analysis in organoids, we used the pySCENIC (v0.11.2) package.^7^ The Loom file of each organoid dataset was used, and the regulon-based UMAP was redrawn after binarization of each cell with regulons and an AUCell calculation. The AUCell of each regulon and cell cluster were combined to obtain a regulon specificity score, and we found the top regulons of each cluster. These processes were repeated five times in each organoid dataset (WT, PC, PN, and PCN). To find the Ccl2 relevance, we created modules of candidate transcription factors (Rela, Foxa1, Myc, Taf1, and Tcf12) in the PCN dataset. Transcription factors and their target genes were calculated and visualized using Cytoscape.

**Cell-to-cell interaction analysis.** CellChat (v1.4.0) and Squidpy (v1.1.2) packages were used for the cell-to-cell interaction inference. Epithelial cells, fibroblasts, and immune cell datasets were merged at each disease stage to generate gene expression matrices for the CellChat analysis. The merged Seurat objects were converted to an H5ad file for the Squidpy analysis. In the CellChat, the “CCL” pathway was specified for analysis. “Epithelial cell” was defined as a “source group” and “T cell”, “myeloid cell”, and “B cell” were selected as “target groups” in the Squidpy analysis (permutation = 100, *P* value threshold = 0.05).

**Pathway score analysis.** We used Scanpy with the “scanpy.tl.score_genes” function for the pathway score analysis. The analysis was performed with default parameters and the reference genes from the gene ontology biological process or the Kyoto Encyclopedia of Genes and Genomes database. The gene list for the score analysis is shown in Supplementary Table 8.

**Chromatin immunoprecipitation assay**

A chromatin immunoprecipitation (ChIP) assay was performed as previously described.^8^ In brief, PCN cells were harvested after being crosslinked with 1% formaldehyde for 10 min at ambient temperature, followed by quenching with glycine (final concentration = 0.125M). Cells were lysed and subjected to sonication (15 rounds of 30 sec on and 30 sec off, Bioruptor 300 [Diagenode]). Lysates were cleared and incubated with pre-conjugated RELA antibody-Dynabeads overnight at 4 °C. Immunoprecipitates were washed and eluted, followed by incubation with RNase. After RELA protein degradation with Proteinase K, ChIP DNA was purified using a PCR purification kit (QIAGEN, #28106). Putative Rela binding sites of the *Ccl2* promoter regions (region 1: -673~-683, region 2: -1803~-1813, and region 3: -2035~-2045) and non-specific regions (-9708~-9838; a negative control) were subjected to ChIP-qPCR. The primers for ChIP-qPCR are listed in Supplementary Table 2.

**Kaplan-Meier analysis**

The overall survival of ESCC patients based on gene expression was determined using a publicly available database (Kaplan-Meier Plotter, <http://kmplot.com/analysis>). Eighty-one ESCC patients were divided into “high” and “low” groups by the median expression of the gene of interest. The overall survival of the patient groups was compared for 70 months, and the log-rank *P* value is marked in the figure.

**Quantification and statistical analysis**

The Student’s *t*-test was used for comparisons of two groups (n ≥ 3), and one-way analysis of statistical variance evaluation was used for comparisons of at least three groups (n ≥ 3). *P* values < 0.05 were considered significant. Error bars indicate the standard deviation (s.d.). All experiments were performed three or more times independently under identical or similar conditions.
