## Supplementary information for "Key Genetic Determinants Driving Esophageal Squamous Cell Carcinoma Initiation and Immune Evasion"

**Supplemental Information**

**Supplementary Figures**

Supplementary Figure S1. Candidate genes KO in EOs.

Supplementary Figure S2. Establishment of CRISPR/Cas9-based KO organoids and validation.

Supplementary Figure S3. *Notch1* KO suppresses organoid differentiation and induces cell migration.

Supplementary Figure S4. Integration and annotation of scRNA datasets of organoids.

Supplementary Figure S5. CCL2 relevance in immune response.

Supplementary Figure S6. TME comparison of PCN- and PCN-*Ccl2* KO–transplanted tumors.

Supplementary Figure S7. Unsupervised clustering and annotation of epithelial cells and immune cells of mouse ESCC model datasets.

Supplementary Figure S8. CCL2-CCR2 interaction in epithelial and immune cells of the mouse ESCC model.

Supplementary Figure S9. Immune landscape regulation by CCR2 inhibitors during tumorigenesis and clinical evidence of Rela/NF-κB-CCL2 axis.

Supplementary Figure S10. Characterization of ESCC patient subgroups.

**Supplementary** **Figure S1. Candidate genes KO in EOs.**

**A,** Oncogrids of nine candidate genes were generated based on the mutation and copy number alterations. Eighty-six patients with ESCC were analyzed from the TCGA database.

**B,** Mutation types of candidate genes were analyzed in the patients’ data from cBioportal and the TCGA. The genes harboring more than 50% of truncation or frameshift mutations are shown.

**C,** Gene expression of FAT4 and KMT2C from ESCC patients (TCGA, *n* = 95) and normal (TCGA, *n* = 11) samples are shown with box plots. Data are shown as means ± SEM.

**D,** Genetic status of each organoid line was designed as shown in the table.

**E,** Bright-field images of organoids are shown. Images were taken on day 8 of passage 3. Scale bar = 50 μm.

**F,** Hematoxylin-and-eosin-stained sections from KO organoids. Scale bar = 50 μm.

**Supplementary** **Figure S2. Establishment of CRISPR/Cas9-based KO organoids and validation.**

**A,** Representative images of mouse EO growth of *Trp53^floxed/floxed^* are shown. Images were taken from day 3 to day 9 of passage 3. Scale bar = 20 μm.

**B,** KO efficiency of each sgRNA was validated with immunoblot. Three different sgRNA constructs of each gene were tested with transient transfection.

**C,** Genomic DNA of each organoid was sequenced to verify CRISPR/Cas9-induced gene editing. Pink boxes show the sgRNA sequences of each gene.

**Supplementary** **Figure S3**. ***Notch1* KO suppresses organoid differentiation and induces cell migration.**

**A-B,** Organoids were stained with MKI67, SOX2 **(A)**, and KRT13 **(B)**. PA: *Trp53^del/del^* + *Ajuba* KO, PCF4: *Trp53^del/del^* + *Cdkn2a* KO + *Fat4* KO, PCA: *Trp53^del/del^* + *Cdkn2a* KO + *Ajuba* KO, WT: *Trp53^floxed/floxed^*, N: *Notch1* KO, C: *Cdkn2a* KO, CN: *Cdkn2a* KO + *Notch1* KO, and P: *Trp53^del/del^*. Scale bar = 20 μm.

**C,** Morphologies and internal structures of DAPT-treated P and PC organoids were compared with PN and PCN, respectively. DAPT was treated with different doses; low dose = 5 mM, high dose = 10 mM. Scale bar = 50 μm.

**D,** A reversible Notch1 inhibition effect was shown in the bright-field microscopic images. Morphologies of P and PC were rescued after DAPT removal in the second passage. DAPT (10 mM) was treated for 8 days for each passage. Upper panel, magnification, ×50, Scale bar = 200 μm. Lower panel, magnification, ×100, Scale bar = 100 μm.

**E,** Schematic overview of organotypic culture experiment.

**F,** Bright-field images of 2D-cultured cells. Images were taken on day 3 of passage 3. Scale bar = 100 μm.

**G,** Wound closure rates of PC, PN, and PCN cells were evaluated by analyzing images taken at 0, 8, and 20 hrs after scratch with ImageJ software.

**H,** 2D-cultured PCA, PCF4, PN, PCN cells at day 3 of first and second passages.

**I,** Alcian Blue-PAS staining results of organoids are shown. The large intestine tissue was used as a positive control. Scale bar = 50 μm.

**Supplementary** **Figure S4. Integration and annotation of scRNA datasets** **of organoids.**

**A,** Integrated UMAP of WT, PC, PN, and PCN datasets (Seurat package).

**B,** Heatmap of each cluster from the integrated dataset.

**C,** Cells were clustered by 35 subtypes and annotated as four cell types based on the marker genes.

**D,** Marker gene expression of each cell type shown in the dot plot.

**E,** The proportion comparison of each cell cluster from the PC dataset and WT.

**Supplementary** **Figure S5**. **CCL2 relevance in immune response.**

**A-C,** 5-week-old scid-mice were injected with 5x10^6^ cells/mouse of PN and PCN, and tumors were collected 35 days after transplantation (**A**). Tumor growth was assessed by tumor weight (**B**) and volume (**C**).

**D,** H&E images of PN and PCN-derived tumors from scid-mice. Scale bars = 100 μm (left panels) and 20 μm (right panels).

**E,** SOX2 staining of PN (top) and PCN (bottom) tumors from scid-mice. Scale bars = 100 μm (left panels) and 20 μm (right panels).

**F,** Tumor-infiltrated immune cells of PCN-transplanted tumor tissues were stained with Havcr2/Tim3, Pdcd1/Pd-1, Cd8, Cd4, Mpo, and Cd209/Dc-sign. Nuclei were stained with DAPI. Scale bar = 50 μm. Dotted circle, inflammatory cells.

**G,** An Enrichr analysis from PCN highly expressed genes in proliferating cells from the scRNA-seq results of PN and PCN datasets. DEGs were created between proliferating cells of PCN and PN, and the relevant pathways were analyzed with BioPlanet and the KEGG databases.

**H,** An Enrichr analysis with bulk RNA-seq results. PCN highly expressed genes compared to PN were analyzed using gene ontology biological process (GOBP) database.

**I,** Dot plots showing the gene expression of *Ccl2*, *Cxcl1*, and *Cxcl2* in a different cell type and datasets of organoids from scRNA-seq results.

**J,** *Ccl2* expression in the PC dataset displayed on UMAP.

**K,** Dot plot from scRNA-seq showing *Ccl2* gene expression in different genotypes of organoids.

**L-M,** *Cxcl1* and *Cxcl2* expression in different genotypes of organoids shown on a dot plot (**L**) and UMAP (**M**). **N,** PCN-transplanted tumors were stained with CCL2 and CCR2 antibodies. Scale bar = 50 μm.

**Supplementary** **Figure S6. TME comparison of PCN- and PCN-*Ccl2* KO–transplanted tumors.**

**A,** Sanger sequencing results of PCN and PCN-*Ccl2* KO cells. Pink box showing the sgRNA sequences for the *Ccl2* gene.

**B,** Growth rates of PCN and PCN-*Ccl2* KO cells.

**C-D,** 5-week-old scid-mice were injected with 5x10^6^ cells/mouse of PCN and PCN-*Ccl2* KO cells, and tumors were collected 35 days after transplantation. Tumor growth was assessed by tumor weight (**C**) and volume (**D**).

**E,** H&E staining of PCN- and PCN-*Ccl2* KO–derived tumors from immunocompetent mice. Scale bar = 100 μm.

**D,** PDCD1/PD-1, CD8, CD3, CD206, CD209, CD68, CD11B, LY6G, cleaved-Caspase3 (cCas3), and MKI67 staining in PCN- and PCN-*Ccl2* KO cell–derived tumors from immunocompetent mice. Samples were counterstained with DAPI. Scale bar = 100 μm.

**E,** Randomly chosen 630× magnified images were analyzed and plotted. n.s. = not significant.

**F,** Higher magnification (630×) images of CD68, CD80, CD11B, and LY6G staining. Scale bar = 20 μm.

**Supplementary** **Figure S7.** **Unsupervised clustering and annotation of epithelial cells and immune cells of mouse ESCC model datasets.**

**A,** Heatmap showing the marker genes of each cluster of epithelial cells.

**B,** scRNA-seq data of 4-NQO-treated mouse esophagus epithelial cells were projected with UMAP and clustered by disease status. Normal: 0 week of 4-NQO treatment, inflammation: 12 weeks of 4-NQO treatment, hyperplasia: 20 weeks of 4-NQO treatment, dysplasia: 22 weeks of 4-NQO treatment, cancer *in situ*: 24 weeks of 4-NQO treatment, invasive cancer: 26 weeks of 4-NQO treatment.

**C,** UMAP projection of cells with cell type annotations based on the marker gene expression.

**D,** Heatmap showing the marker genes of the immune cell sub-clusters.

**E,** Immune cells of each disease status are clustered by cell types and displayed with UMAP.

**F,** Detailed sub-clusters of immune cells are annotated by marker gene expression using the CellKb database.

**Supplementary** **Figure S8**. **CCL2-CCR2 interaction in epithelial and immune cells of the mouse ESCC model.**

**A,** The expression of Trp53, Cdkn2a, and Notch signaling pathways-related genes were displayed at the different disease stages after 4-NQO treatment by dot plot.

**B,** ESCC patients-associated cells were assessed using the Scissor package and displayed with red dots in the UMAP. Blue dots represent normal cell-associated cells, and gray dots are background cells.

**C,** ESCC-associated cell proportions of each disease status are displayed with bar plots.

**D-E,** Feature plot **(D)** and dot plot **(E)** of *Ccr2* expression in immune cells. Plots were visualized on the basis of the disease status.

**F,** *Ccr2* expression displayed in the dot plot for the sub-clusters of T cells.

**G,** T_ex_ cell marker gene expression in the immune cell clusters were displayed with a dot plot.

**H,** Predicting ligand-receptor interactions between epithelial and immune cells at different disease stages (Squidpy package).

**I,** Ligand-receptor interactions related to the CCL pathway were predicted in sub-clusters of immune cells and epithelial cells. Interactions in the CCL pathway in each disease status (inflammation, dysplasia, and invasive cancer) were calculated and visualized with the circle plot.

**J,** Significant interactions of the CCL pathway in epithelial cells with T_ex_ cell, MDSC, and macrophage sub-clusters were plotted in inflammation, dysplasia, and invasive cancer status using a chord diagram. The predicted genes were annotated for ligands in epithelial cells and receptors in immune cells.

**Supplementary** **Figure S9. Immune landscape regulation by CCR2 inhibitors during tumorigenesis** **and clinical evidence of Rela/NF-κB-CCL2 axis.**

**A,** qRT-PCR results showing the relative mRNA expression of *Ccl2* in PCN and PCN ectopically expressing Sox2 (PCNS) cells. n.s. = not significant.

**B-C,** Tumorigenicity comparison in PCN and PCNS cells.

**D,** Cell growth of PCNS was assessed after treatment with DMSO or 1 μM of CCR2 inhibitors (CCR2 22 and BMS-813160) by measuring the cell number.

**E,** H&E staining of vehicle-, CCR2 22-, and BMS-813160-treated tumors.

**F,** PDCD1/PD-1, CD8, MKI67, CD206, CD68, and CD209 staining images in vehicle-, CCR2 22-, and BMS-813160-treated tumors. Scale bar = 100 μm.

**G-H,** Images of high magnification (630×) from CD8 and CD209 staining results in vehicle-, CCR2 22-, and BMS-813160-treated tumors **(G)** and quantitation of CD209^+^ cells **(H)**. Scale bar = 20 μm. ***P* < 0.01, n.s. = not significant.

**I,** Human ESCC tissue microarray slides from 112 samples were stained with anti-RELA and CCL2 antibodies. Scale bar = 50 μm (lower magnification) and 20 μm (higher magnification).

**J,** Correlations between RELA and CCL2 expression are displayed with a heatmap. The Pearson correlation coefficient (r) and *P* value (*p*) are displayed. *n* = number of samples.

**Supplementary** **Figure S10. Characterization of ESCC patient subgroups.**

**A,** *CD274* (*PD-L1*) expression in each patient shown with a dot plot.

**B-C,** Pathway scores specifically enriched in ESC2 (**B**) and ESC3 (**C**) were shown with dot plots.

**D,** Stacked bar plot with each group of patients and their tumor grade.

**E,** Expression of marker genes of each patient group.

**F,** Expression of marker genes from each group was assessed by survival duration. A Kaplan-Meier plot was drawn by splitting ESCC patients from the TCGA database by the median value of each gene expression.

**G,** Expression levels of marker genes of each group were measured in ESCC patients of the TCGA database and shown with box plots. NT: normal tissue of EAC and ESCC, TP: tumor from ESCC patients.

**H,** Heatmap display of the correlation matrix of human ESCC tissue microarray sample staining results of B2M, CCL2, and RELA.
